# spSeudoMap: Cell type mapping of spatial transcriptomics using unmatched single-cell RNA-seq data

**DOI:** 10.1101/2022.05.09.491104

**Authors:** Sungwoo Bae, Hongyoon Choi, Dong Soo Lee

**Affiliations:** Institute of Radiation Medicine, Medical Research Center, Seoul National University, Seoul, Republic of Korea; Department of Nuclear Medicine, Seoul National University Hospital, Seoul, Republic of Korea; Department of Nuclear Medicine, Seoul National University College of Medicine, Republic of Korea; Portrai, Inc. Republic of Korea

**Author notes:** **Correspondence**, Hongyoon Choi, MD., Ph.D. Department of Nuclear Medicine, Seoul National University Hospital, 101, Daehak-ro, Jongno-gu, Seoul 03080, Republic of Korea, Dong Soo Lee, MD., Ph.D. Department of Molecular Medicine and Biopharmaceutical Sciences, Graduate School of Convergence Science and Technology, Seoul National University, 103, Daehak-ro, Jongno-gu, Seoul 03080, Republic of Korea.

**Keywords:** spatial transcriptomics, single-cell RNA-seq, cell sorting, cell type mapping, synthetic cell mixture, pseudobulk

## Abstract

With advances in computational models, the cellular landscape can be tracked in various tissues using spatial transcriptomics. Since many single-cell RNA-seq (scRNA-seq) data have been obtained after cell sorting, such as when investigating immune cells, integrating these singlecell data with spatial data is limited due to a mismatch of cell types composing the two datasets. Here, we present a method, spSeudoMap, which utilizes sorted scRNA-seq data to train a model for predicting cell types of spatial spots by creating virtual cell mixtures that closely mimic the gene expression profile of spatial transcriptomic data. To overcome the mismatch issue, the cell type exclusively present in the spatial data, pseudotype, was defined. The proportion of pseudotype cells and virtual expression profiles in the cell mixture was determined by pseudobulk transcriptomes. The simulated cell mixture was considered a reference dataset, and the model that predicts the cell composition of the mixture was trained to predict the cell fraction of the spatial data using domain adaptation. First, spSeudoMap was evaluated in human and mouse brain tissues, and the main region-specific neuron types extracted from single-cell data could be precisely mapped to the expected anatomical locations. Moreover, the method was applied to human breast cancer data and described the spatial distribution of immune cell subtypes and their interactions in heterogeneous tissue. Taken together, spSeudoMap is a platform that predicts the spatial composition of cell subpopulations using sorted scRNA-seq data, and it may help to clarify the roles of a few but crucial cell types.

## Main

Spatial transcriptomics has been widely adopted as a tool to explore genome-wide spatial RNA expression in various tissues^1^. It paves the way to thoroughly investigate the spatial context of cells and their interactions in an unbiased manner^2^. One of the limitations of spatial transcriptomics data is the fact that spots are not directly interpreted as cells. Therefore, multiple computational approaches have been suggested for accurate spatial mapping of cell types by integrating spatial and single-cell transcriptomics^3–10^. They can be further utilized to speculate the spatial infiltration pattern of a few cell types that play a key role in the pathophysiology of various diseases^11–14^. In this case, certain cell populations, such as immune cell subtypes, can be described in detail by jointly analyzing spatial data with single-cell RNA sequencing (scRNA-seq) data acquired from cell sorting strategies based on cell surface markers^15^. However, there is a major drawback in the practical usage of spatial mapping methods employing single-cell RNA-seq data. The majority of the methods are based on the assumption that cell types are similar between the two transcriptomes^4–10^, and cell type-specific signatures defined from the single-cell data are translated to decipher spatial cell compositions. Therefore, when the sorted scRNA-seq data are utilized as a reference, the data explain only part of the cell types from the spatial data, and the integration methods create bias in estimating the cellular fraction. There is a need for the development of a computational model that flexibly integrates scRNA-seq data of cell subpopulations with spatial transcriptomic data.

In this regard, we suggested a model, spSeudoMap, which utilizes sorted scRNA-seq data to create a cell mixture resembling the spatial data and predicts the spatial cell composition based on CellDART^9^, a domain adaptation approach. More specifically, the cell type exclusively present in the spatial data, named the pseudotype, was defined. Then, the fraction and expression profiles of the pseudotype in the cell mixture were assigned by referencing the spatial and single-cell transcriptomes. The rest of the mixture was filled with randomly sampled cells from single-cell data. As a result, the modified cell mixture and spatial transcriptomic data were jointly analyzed to obtain a spatial map of the cell subpopulation.

## Results

### Mapping the excitatory neuron subpopulation in the human brain with spSeudoMap

The capability of the method, spSeudoMap, was assessed in the Visium spatial transcriptomes of the human dorsolateral prefrontal cortex (DLPFC). First, 10 layer-specific excitatory neuron subtypes were selected from the single-cell data of the human brain (for convenience, single-nucleus RNA-seq data were also called single-cell data) (**Supplementary Fig. 1**), and the cell subpopulation was jointly analyzed with the spatial data. Notably, the physical sorting of excitatory neurons did not precede scRNA-seq. Instead, for simulation purposes, excitatory neuron subpopulations were manually selected from the whole scRNA-seq data, and the cell types were mapped to the spatial transcriptomic data. A pseudospot, the cell mixture that contains all cell types in spatial data, was defined, and the proportion and expression of the ‘pseudotype’ were designated based on both transcriptomes (**Fig. 1**). First, pseudobulk gene expression profiles were computed from both datasets. The top genes highly expressed in spatial transcriptomic data compared to single-cell data were selected as virtual pseudotype markers. These markers represented cell types that were not included in scRNA-seq data. Next, a pseudotype fraction in spatial spots was estimated by calculating module scores of pseudotype markers. Then, the pseudotype proportion and expression profiles in the cell mixture were designated by referencing the presumed pseudotype fraction and expression in a randomly selected spatial spot. The remaining portion of the mixture was filled with randomly sampled cells from single-cell data. Finally, the modified pseudospot was considered a reference dataset for the domain adaptation model CellDART^9^.

**Fig. 1.**
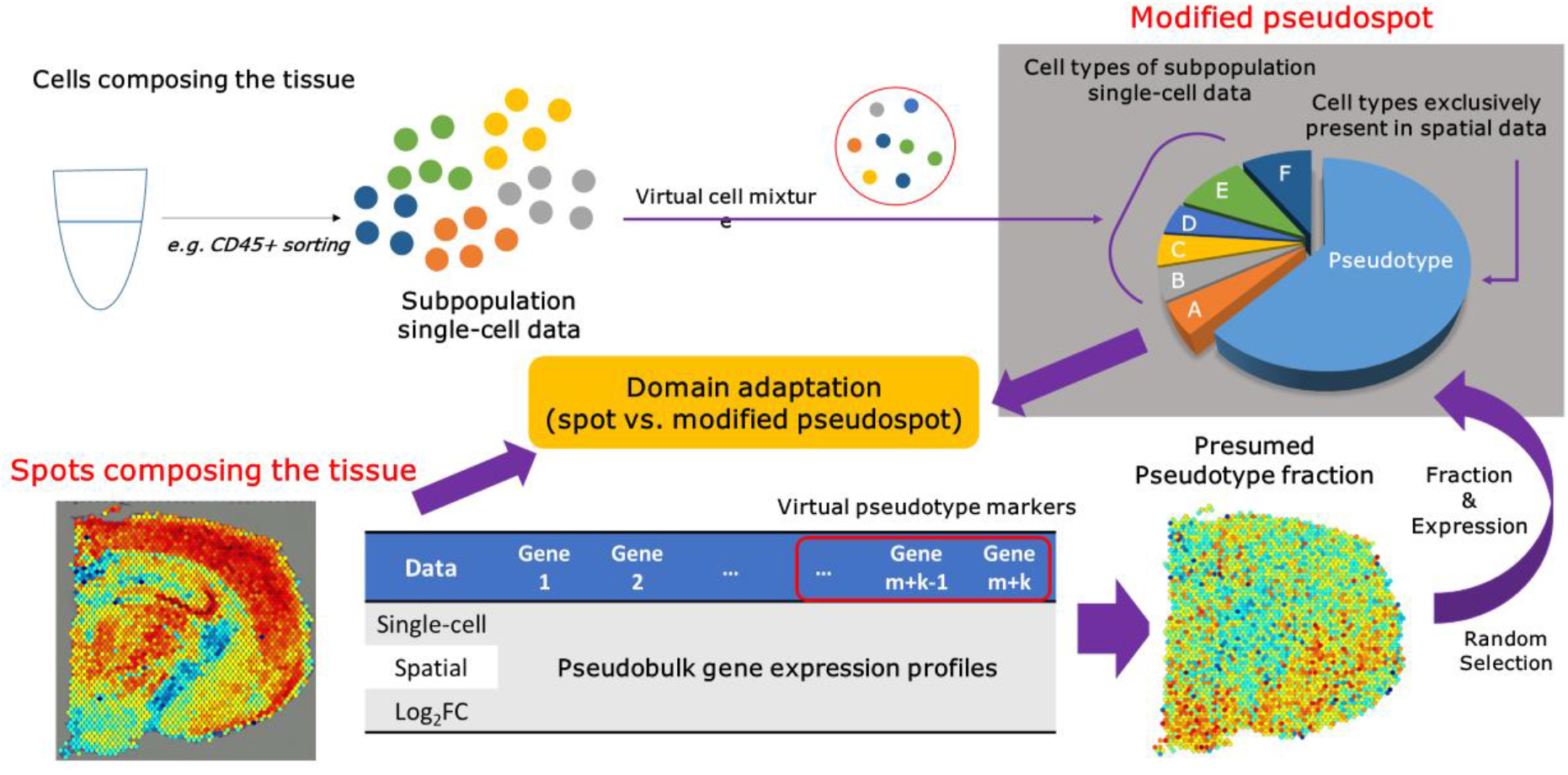
Mapping of cell subtypes to the spatial transcriptome with spSeudoMap. The cell types of the subpopulation single-cell transcriptome acquired from cell sorting experiments can be spatially mapped to the tissue using spSeudoMap. The subpopulation single-cell data are composed of the sorted cells from the tissue, and the cell types cover only part of those in the spatial transcriptome. To create the reference dataset that mimics the spatial data, virtual cell mixtures were defined in which all cell types from the tissue were included. First, the cell types exclusively present in the spatial data were aggregated and named pseudotype. The virtual markers for the pseudotype were selected from the top genes highly expressed in spatial pseudobulk compared to single-cell pseudobulk data. Then, the pseudotype fraction in the spatial spots was estimated from the module scores (sc.tl.score_genes in Scanpy) of the top 20 pseudotype markers. The fraction and gene expression of the pseudotype were assigned based on the presumed pseudotype fraction and expression of a randomly selected spatial spot. Last, the nonpseudotype proportion of the modified pseudospot was filled with the randomly sampled cells from the subpopulation single-cell data. Finally, the modified pseudospot was considered as a reference dataset for the domain adaptation method CellDART^9^.

The optimal parameters for spSeudoMap were searched on the human DLPFC slide (slide no. 151676) (**Methods and Supplementary Fig. 2**). The parameters included the number of markers (*N_markers*), the ratio of the number of single-cell markers to pseudotype markers (*m/k* ratio), and the standard deviation (SD) of the pseudotype fraction (*Nmarkers=80, m/k* ratio=4, and *pseudof_std =* 0.1). The layer-specific excitatory neuron fraction predicted by the model was mapped to the tissue. Overall, the cell types were highly localized to the corresponding cortical layers (e.g., layers 4 to 5 for Ex_3_L4_5), although Ex_1_L5_6 and Ex_8_5_6 showed uneven distribution patterns within the same layer (**Fig. 2a**).

**Fig. 2.**
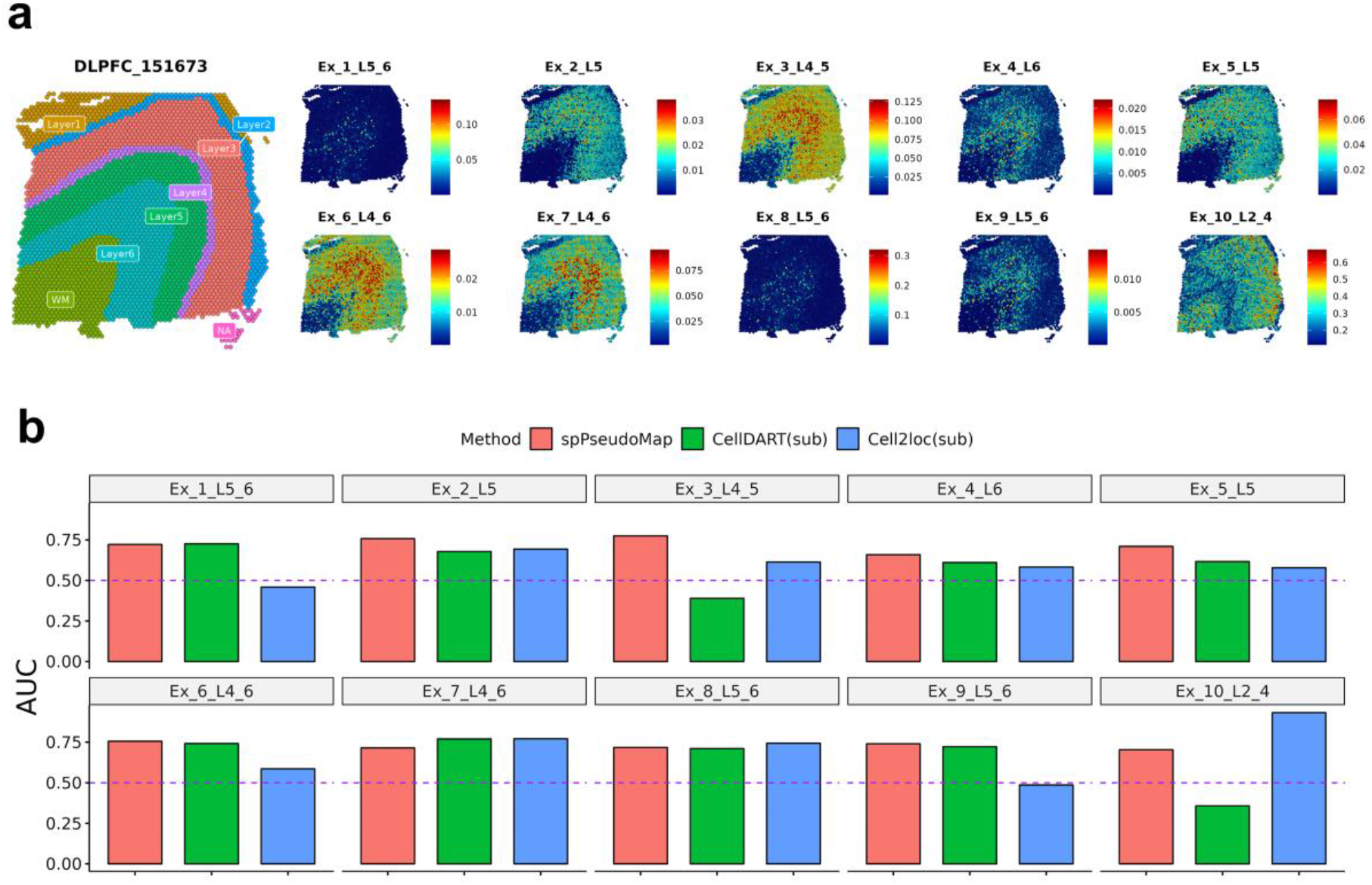
Decoding spatial maps of layer-specific neurons in the human brain. **a**, The composition of ten excitatory neurons in the human brain was predicted by spSeudoMap. The estimated cell fraction was spatially mapped to the tissue and visualized with colored bars. Overall, the spatial distribution of layer-specific neuron types was restricted to the corresponding cortical layer (from layers 1 to 6). **b**, The performance of the computational models was evaluated by layer discriminative accuracy of the predicted neuron fraction and represented by the area under the receiver operating characteristic curve (AUROC). The AUC values of spSeudoMap in 10 cell subtypes were visualized with barplots and compared with those of CellDART and Cell2location. Both CellDART and Cell2location were implemented for the subpopulation single-cell data. SpSeudoMap was overall superior to CellDART except for Ex_1_L5_6 and Ex_7_L4_6 and superior to Cell2location except for Ex_7_L4_6, Ex_8_L5_6, and Ex_10_L2_4.

### Comparison with cell type mapping tools using subpopulations in the human brain

For the next step, the subpopulation single-cell data for 10 neuron types and spatial data were integrated with CellDART and Cell2location tools, and the spatial composition of the cell types was predicted. It was intended to evaluate the performance of the previous integration methods when there was a mismatch of cell types between both transcriptomes. In the case of CellDART, Ex_3_L4_5 and Ex_10_L2_5 had lower cell fractions in the corresponding layers than in other layers (**Supplementary Fig. 3a**). Additionally, in Cell2location, Ex_1_L5_6 and Ex_9_L5_6 had opposite patterns of distribution, with higher cellular abundance in layers other than 5 and 6 (**Supplementary Fig. 3b**).

To measure the layer discriminative accuracy of the predicted cell fraction, an area under the receiver operating characteristic curve (AUROC) was calculated, considering layer annotation as a reference. The AUROC was compared across the three methods (**Fig. 2b**), and spSeudoMap showed superior performance than CellDART in 8 out of 10 and Cell2location in 7 out of 10 cell types. In summary, within the optimal parameter range, spSeudoMap presented more stable performance across all cell types in the neuron subpopulation compared to CellDART and Cell2location.

### Deciphering the excitatory neuron composition in the mouse brain with spSeudoMap

The performance of spSeudoMap was evaluated in coronal sections of the mouse brain (**Supplementary Fig. 4**). First, the two cell type deconvolution tools, CellDART and Cell2location, were tested on the original single-cell data containing all cell types (**Supplementary Fig. 5a**). The two prediction results were considered reference standards for assessing spSeudoMap. In both methods, the region-specific neurons showed spatially restricted patterns according to their expected spatial predominance (**Supplementary Fig. 6**).

The excitatory neuron subpopulation was selected as a simulation of sorted subpopulation scRNA-seq data (**Supplementary Fig. 5b)**and spatially mapped to the brain with spSeudoMap (**Fig. 3a**). The 8 cell types showed a highly restricted distribution with a similar spatial predominance as the reference results (**Fig. 3b and Supplementary Fig. 7**). Other cell types were also mapped on the mouse coronal section data and compared with the reference results (**Supplementary Fig. 8**). Some of the neuron types having a low fraction (<0.05) estimated by spSeudoMap were distributed not only in the expected regions but also in the regions outside of interest. According to the results, spSeudoMap was capable of precisely predicting the spatial compositions of the main cell types composing the cell subpopulation.

**Fig. 3.**
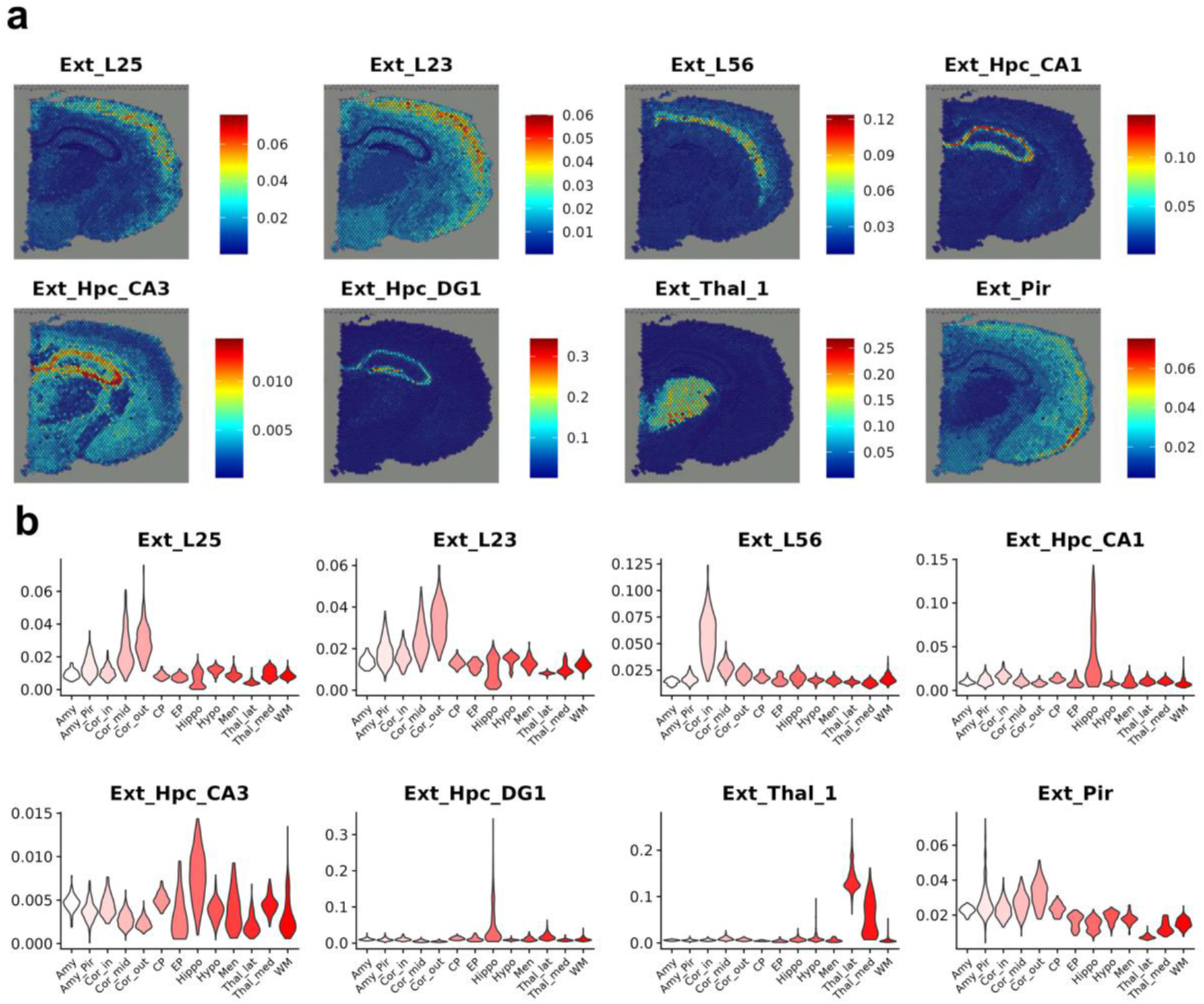
Predicting the spatial composition of region-specific neuron types in the mouse brain. **a**, The proportion of the representative region-specific neuron types was estimated by spSeudoMap and mapped to the mouse brain. Overall, the 8 neuron types presented were predominantly localized to the corresponding anatomical locations. **b**, Spatial spots were clustered based on their gene expression profiles, and the spot clusters were named after the anatomical locations. The predicted neuron fraction was highly distributed according to anatomical location. Ext_L25 to the mid-cortical layer, Ext_23 to the outer cortical layer, Ext_L56 to the inner cortical layer, Ext_Hpc_CA1, Ext_Hpc_CA3, and Ext_Hpc_DG1 to the hippocampus, Ext_Thal_1 to the thalamus, and Ext_Pir to the amygdala or piriform cortex area. Amy: amygdala, Amy_Pir: amygdala or piriform cortex, Cor_out: outer cortex, Cor_mid: mid cortex, Cor_in: inner cortex, CP: caudoputamen, EP: ependyma, Hippo: hippocampus, Hypo: hypothalamus, Men: meninges, Thal_lat: lateral thalamus, Thal_med: medial thalamus, and WM: white matter.

### Elucidating cellular heterogeneity in human breast cancer with spSeudoMap

spSeudoMap was assessed in breast cancer tissue, which has a high level of heterogeneity. We mapped cell types of spatial transcriptomics of human breast cancer using two different scRNA-seq datasets: one dataset with unsorted whole-cell types and another dataset with CD45+ sorted data. The prediction results of CellDART obtained using all cell types in the unsorted single-cell data were considered a reference (**Supplementary Fig. 9a**). Immune cell types from CD45+ sorted single-cell data (**Supplementary Fig. 9b**) were spatially mapped to human breast cancer tissue using spSeudoMap. For macrophages, which are the immune cell type with the highest proportion in the tissue (**Supplementary Fig. 9b**), the spatial cell compositions were similar between CellDART and spSeudoMap (**Fig. 4a, b**). Additionally, CD4+ T cells and B cells showed similar spatial patterns with positive spatial correlation (**Fig. 4c**), although B cells in the two single-cell datasets indicated different cell subtypes. However, the spatial localization patterns of CD8+ NKT-like cells, one of the minor cell types in scRNA-seq data, were different between the two methods. Finally, to investigate the spatial interaction between the immune cells, the pairwise correlations between the predicted cell fractions were computed and visualized with a heatmap (**Supplementary Fig. 10**). Spatial correlation patterns were highly similar between one of the macrophage subtypes and memory CD4+ T cells. Additionally, high proximities were observed between another subtype of macrophages and CD8+ NKT-like cells. These findings implicate the spatial interaction of macrophages with T cells in the tumor tissue^16^. In short, spSeudoMap can be applied to decipher complex tumor microenvironments and to understand spatial interactions between cell subpopulations.

**Fig. 4.**
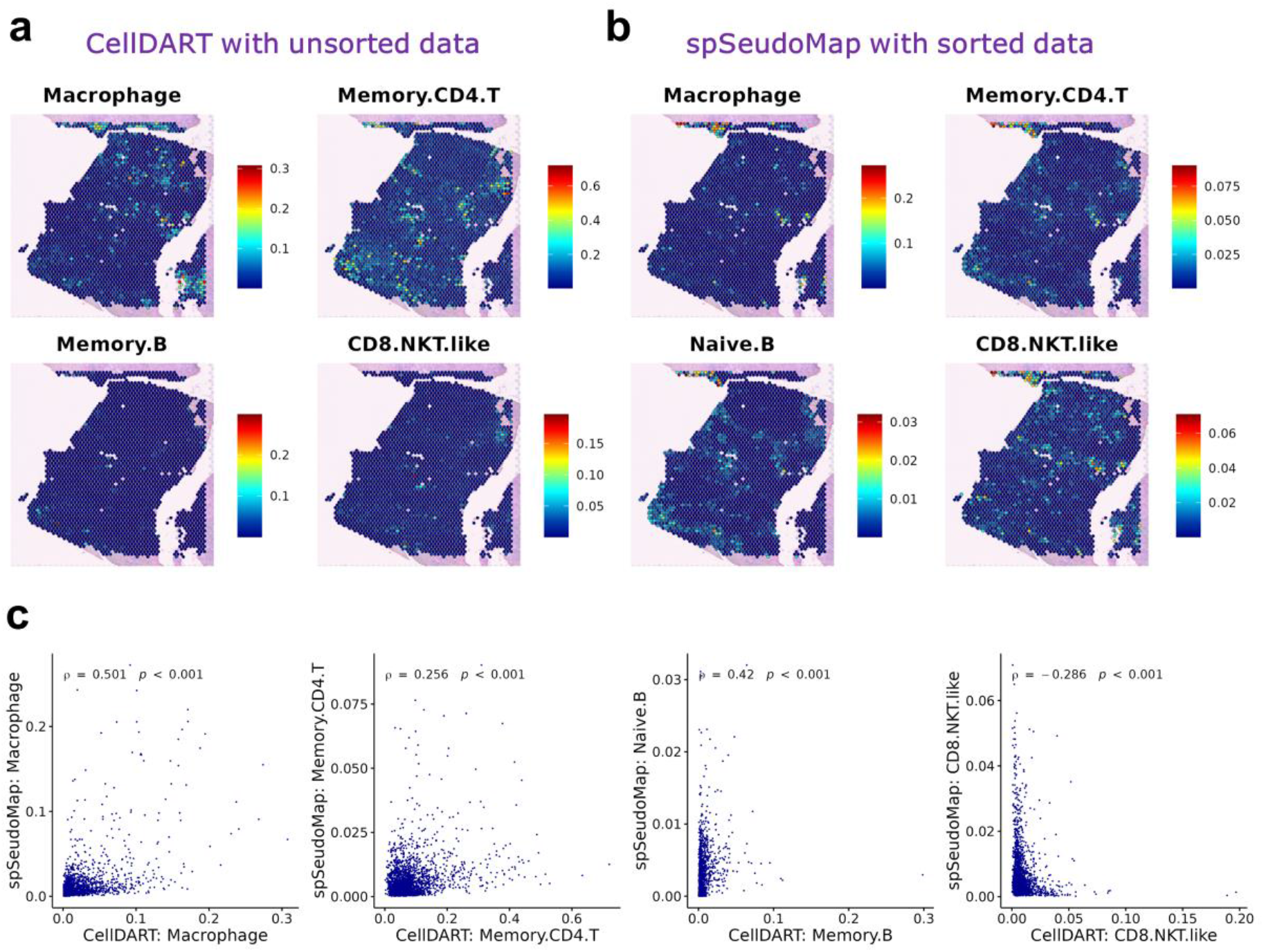
Exploring the spatial heterogeneity of immune cells in human breast cancer tissue. **a**, The spatial composition of immune cells in human breast cancer was predicted by integrating spatial data with single-cell data covering all cell types. The results from CellDART were considered a reference for comparison with spSeudoMap. **b**, The spatial composition of the immune cell types was predicted by spSeudoMap using CD45+ sorted single-cell data and visualized on the tissue. Macrophages (sum of macrophage_1, macrophage_2, and macrophage_3), memory CD4 T cells, and B cells showed similar spatial distributions, while CD8 NKT-like cells presented different patterns. **c**, The scatter plots show the correlation between the cellular proportion predicted by spSeudoMap with sorted single-cell data and that estimated by CellDART with an unsorted single-cell dataset covering all cell types. Spearman’s correlation coefficients and statistical significance (p-value) were calculated and are presented in the top-left corner of each plot. The correlation between CellDART and spSeudoMap was the highest in macrophages (the sum of macrophage_1, macrophage_2, and macrophage_3). Other cell types also showed weak but positive correlations except for CD8+ NKT-like cells, which showed a negative correlation.

## Discussion

Exploring the spatial composition of infiltrating cells in tissues provides a key to understanding the molecular mechanism underlying the functional changes. In recent years, the crosstalk between immune cells and cancer cells has been highlighted as a major player in the development of tumor microenvironments^12^. Additionally, the complex interactions between the peripheral and central immunities occurring at brain borders are considered crucial for the progression of various neuroinflammatory diseases^13^. As a breakthrough to the research questions, the integration of spatial transcriptomics with the subpopulation single-cell data can provide a cellular landscape of the few cell types, such as immune cells, in highly heterogeneous tissues.

Compared to the existing approaches, spSeudoMap, which is specialized in tracking the spatial distribution of subclusters of specific cell types, is more appropriate for capturing the transcriptomic changes in a few cell populations. The key contribution is the integration of scRNA-seq of specific cell subtypes into spatial transcriptomics. First, spSeudoMap was assessed in human and mouse brain tissues, and the main region-specific excitatory neuron types could be stably mapped according to anatomical locations (**Fig. 2, 3**). Next, it could be extended to human breast cancer tissue to estimate the spatial distribution of immune cell types (**Fig. 4**). The predicted spatial patterns of the major immune cell populations were well correlated with the reference results, in which original single-cell data with all cell types were utilized for integration.

One of the key features of spSeudoMap is that it assumes the virtual cell type, pseudotype, which is absent in single-cell data but present in the spatial data. Then, it finds markers of the pseudotype by considering single-cell and spatial data as bulk sequencing datasets (**Fig. 1**). Moreover, spSeudoMap predicts the pseudotype fraction in each tissue domain and adopts the information to model the reference dataset, modified pseudospots. These processes allow the cell mixture to be closer to the spatial transcriptome in terms of gene expression. This is supported by the result that the presumed pseudotype fraction that was utilized for the generation of the modified pseudospot correlates with the pseudotype fraction from the output of spSeudoMap (**Supplementary Fig. 11**). In contrast, the existing methods utilize cell type signatures solely extracted from the subpopulation single-cell datasets, and it will be biased to select the genes differentially expressed inside of the cell subset. Thus, the signatures would not well represent the real expression profiles of cell types in the tissue. This may result in an imprecise estimation of the cellular composition. In this regard, some layer-specific excitatory neuron types showed nonspecific distribution when the subpopulation single-cell data were directly given as inputs to CellDART and Cell2location (**Supplementary Fig. 3**).

As in the aforementioned simulation examples (**Fig. 2, 3**), spSeudoMap can be applied to investigate the spatial compositions of the cell subtypes of a certain cluster. When the original single-cell data are composed of finely defined clusters and the clusters are closely located in terms of gene expression, their marker genes will largely overlap. It may interfere with the precise computation of cell fractions with the existing deconvolution tools. In that case, spSeudoMap can be applied as an alternative, and the selected cell subtypes and their count matrix are provided as inputs to the model. Furthermore, spSeudoMap can be directly applied to enriched single-cell data in which certain cell types are collected from multiple samples by cell sorting strategies. The cell types without enrichment can be removed, and the remaining single-cell data can be used to create the spatial map of the enriched cell types.

There are additional considerations when implementing spSeudoMap to predict the spatial composition of the cell subpopulation. Among the main parameters of the model, the mean of the pseudotype fraction (*pseudofrac_m*) must be given to generate modified pseudospots. For the single-cell data obtained from the cell sorting experiments, *pseudofrac_m* can be considered a fraction of the negative population during the sorting. Alternatively, *pseudofrac_m* can be determined based on the literature evidence that explains the cell type proportion. In fact, spSeudoMap is a generalized form of CellDART, and it flexibly models various situations. If the single-cell and spatial transcriptomes have highly similar cell types, the mean and standard deviation of the pseudotype fraction can be set to 0, and then spSeudoMap is identical to CellDART. When both datasets have significantly different cell types, only the shared cell types can be selected, and their proportions and expression profiles can be given as inputs. In that case, *pseudofrac_m* will be closer to 1 than 0. Meanwhile, in spSeudoMap, the distribution of the pseudotype fraction is assumed to be linearly proportional to the gene set enrichment score of the top pseudotype markers. Although the real distribution may be different, the domain adaptation process in CellDART could have managed the discrepancy between the modified pseudospots and spatial datasets. Last, when the rare cell types are mapped to the tissue, the cellular fraction may be overestimated in the regions outside those of interest, and the region-specific localization patterns may be less prominent. In addition, when the single-cell data contain unmatched cell types compared to the spatial data, as in the basophils and effector CD8 T cells in sorted single-cell data of breast cancer (**Supplementary Fig. 10b**), the prediction of minor immune cell fraction may be affected. Thus, caution is required when interpreting the results for the rare cell population.

In summary, spSeudoMap is a robust model to estimate the spatial configuration of cell subtypes explained by sorted scRNA-seq data in tissues with a high level of heterogeneity. It can be further utilized to describe the perturbation of complex intercellular interactions during disease progression, therefore capturing the spatial dynamics of pathophysiology.

## Methods

### Composition of public datasets

#### Human brain cortex

Spatial transcriptomes for human DLPFC tissues were acquired from subjects without neurological disorder^17^. The Visium slides named ‘151673’ and ‘151676’ were selected, and a count matrix (151673: 33,538 genes across 3,639 spots; 151676: 33,538 genes across 3,460 spots) and cortical layer information [from layer 1 to 6, white matter (WM), and not available (NA)] of spatial spots were utilized. For the integrative analysis, a single-nucleus dataset obtained from the DLPFC of healthy individuals was adopted^18^. The count matrix (30,062 genes across 35,212 cells) and the cell type annotations defined based on the well-known marker genes were used^18^. Among the 33 cell types (**Supplementary Fig. 1a**), 10 layer-specific excitatory neurons (from Ex_1_L5_6 to Ex_10_L2_4) were selected (**Supplementary Fig. 1b**), and the cell subpopulation data were provided as an input for spSeudoMap.

#### Mouse brain coronal section

The Visium spatial transcriptome for the mouse brain coronal section (‘V1_Adult_Mouse_Brain’) was downloaded from the 10X Genomics homepage. Additionally, single-nucleus data extracted from mouse brain coronal tissue were adopted for the joint analysis^10^. The count matrices were composed of 32,285 genes and 2,702 spots for spatial data and 31,053 genes and 40,532 cells for single-cell data. The cell types of single-cell data were originally defined by reported marker genes^19, 20^ and the *in situ* hybridization dataset offered by the Allen Brain Atlas (https://mouse.brain-map.org/). Among the 59 cell types (**Supplementary Fig. 5a**), 23 region-specific neuron types were selected (**Supplementary Fig. 5b**), and the subpopulation single-cell data were utilized for the generation of the cell mixture.

#### Human breast cancer

Both spatial and single-cell transcriptomes for breast cancer were extracted from an 88-year-old female subject (ID: 4290) without a previous history of treatment^21^. The cancer tissue was invasive ductal carcinoma (IDC) with ER-positive (90%, 3+), PR-positive (30%, 2+), and HER2-negative (1+) profiles, and cancer had invaded the adjacent skin and chest wall (pathologic T stage: T4b). The single-cell data (5,789 cells and 29,733 genes) from the same patient were composed of cancer or normal epithelial cells, perivascular-like (PVL) cells, cancer-associated fibroblasts (CAFs), endothelial cells, and immune cells (**Supplementary Fig. 9a**). These unsorted data were considered a reference dataset for the prediction of spatial cell composition. In addition, CD45+ sorted single-cell data (29,900 cells and 31,993 genes) acquired from ER-positive IDC patients (n=4, ID: BC1, BC2, BC4, BC6) were utilized^22^, and the spatial immune cell composition was estimated using spSeudoMap.

#### Clustering and annotation of single-cell and spatial data

The clustering and visualization of transcriptomes were implemented using Seurat (v.4.0.5) in R (v.4.1.1)^23^. First, count normalization was performed, and the total count was set to 10,000 across all cells and spots. Then, the count matrices were natural log-transformed [ln(1 + X)], and the top 2,000 highly variable genes (HVGs) were chosen by standardizing counts with the mean-variance relationship (vst method)^4^. Next, the matrices were regressed against the total count and scaled such that the mean and standard deviation for HVGs were 1 and 0, respectively. After reducing the dimensionality of the dataset using principal component analysis (PCA), the top 30 PCs were selected for the downstream analysis. A shared nearest neighborhood (SNN) graph was constructed based on pairwise cell-cell distances calculated on PC space. The Louvain community detection algorithm^24^ was applied with a resolution of 0.5, and the resulting clusters were visualized with uniform manifold approximation and projection (UMAP) plots^25^.

In most single-cell datasets, the cell annotation information offered by the paper was adopted and visualized. For mouse brain spatial data, the spot clusters were named according to the corresponding anatomical structure (amygdala, caudoputamen, cortex, ependyma, hypothalamus, meninges, piriform cortex, thalamus, and white matter) defined in Allen Brain Reference Atlases (https://atlas.brain-map.org) (**Supplementary Fig. 4**). Meanwhile, in the unsorted human breast single-cell data, the cell annotation was originally obtained from the pooled single-cell data of all patients, and each cell type of one of the patients (ID: 4290) contained only a small number of cells. Therefore, the cells were reannotated based on the Louvain cell clustering results by an automatic cell type assignment tool^26^ and cell type information from the paper^21^. Additionally, the cells from the CD45+ sorted breast dataset were annotated by the automatic cell type annotation method^26^.

#### spSeudoMap: spatial mapping of the cell subpopulation transcriptome

To estimate a spatial map of cell types for subpopulation single-cell data such as sorted or enriched single-cell datasets, a synthetic cell mixture that contains all cell types from spatial data was defined (**Fig. 1**). It was intended to create a reference dataset that is highly similar to the spatial transcriptome. For each mixture, the fraction of the exclusive cell types from the spatial data was assigned, and their synthetic gene expression profiles were created. The rest of the cell mixture was generated from the single-cell data. The process was implemented in Scanpy (v.1.5.1)^27^ and Numpy in Python (v.3.7).

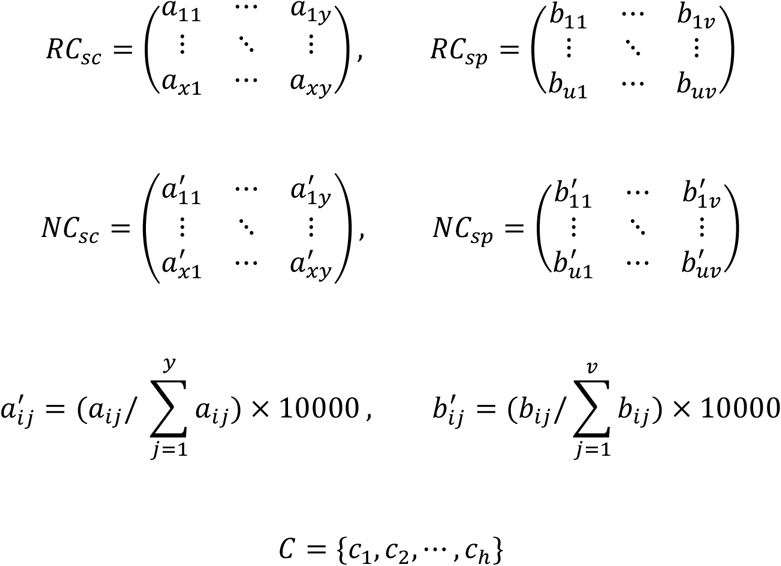

RC_sc_: raw count matrix for single-cell data, RC_sp_: raw count matrix for spatial data, NC_sc_: total normalized single-cell count matrix, NC_sp_: total normalized spatial count matrix, x, y: number of cells and genes in single-cell data, u, v: number of spots and genes in spatial data, C: set of the index for overlapping genes between single-cell and spatial data. The gene index in set C is shared between the two transcriptomes (for example, a_2c_1__ and b_2c_1__ indicate the counts of the same gene with index c_1_ in the second cell and spot, respectively).

First, a fixed number of cells (*n*; brain: 8 and breast: 10) and cell type annotations were randomly sampled from single-cell data with random weights, and a cell mixture named pseudospot was created as with CellDART^9^. The Wilcoxon rank-sum test was performed, and the top *l* markers for each cell type were pooled. A marker panel (total number of genes: *m*) was curated by extracting intersecting genes with the total gene list of spatial data. For each pseudospot, the composite gene expression profiles of the marker panel were calculated.

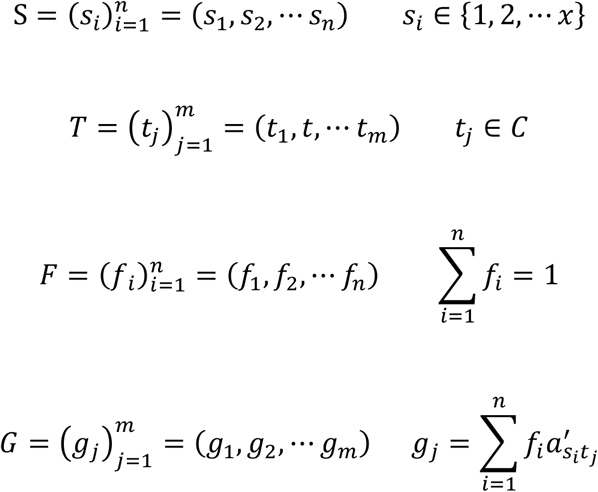

n: total number of the cells to be randomly sampled from the single-cell data, m: total number of marker genes from single-cell data: S: randomly selected index of the cells, T: index of the marker genes in the single-cell count matrix (RC_sc_), F: proportion of each sampled cell in the cell mixture, G: composite gene expression profiles of the cell mixture (named pseudospot)

Next, the cell types present in the spatial data but absent in the single-cell data were aggregated, and the aggregate was named the ‘pseudotype’. The pseudotype markers were speculated from pseudobulk analysis of both transcriptomes. It can be assumed that the genes showing greater pseudobulk expression in spatial data than single-cell data are pseudotype markers. Therefore, the summed counts for each gene were divided by the total counts, and the normalized counts were compared between the two datasets. The ratio of the normalized counts between spatial to single-cell data was log_2_-transformed, and the genes were sorted in descending order of the log fold change. The top *k* genes not overlapping with the single-cell cluster markers were selected as pseudotype markers, and the top 20 genes were used as predictors of the pseudotype fraction. The single-cell and pseudotype markers were combined and named the ‘composite marker panel’.

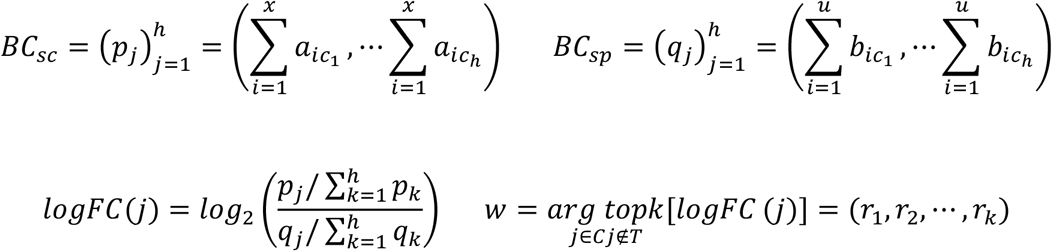

BC_sc_: pseudobulk count matrix for single-cell data, BC_sp_: pseudobulk count matrix for spatial data, logFC: log fold change between the pseudobulk counts, w: index number for the top k genes with the highest logFC

Then, the pseudotype fraction in the spatial data was estimated by calculating an enrichment score for the top 20 pseudotype markers (scanpy.tl.score_genes in Scanpy). The genes of spatial data were divided into 25 bins according to the log-normalized expression level. For each marker gene, a total of 50 control genes were selected from the same bin and pooled. The enrichment score was calculated by subtracting the average expression of control gene pools from that of pseudotype markers^28^. The distribution of a created module score was scaled to have a given mean (*pseudof_m*) and standard deviation (*pseudof_std*). The values over 1 and less than 0 were replaced with 1 and 0, respectively. The scaled module score was considered as a pseudotype fraction of a spatial spot, assuming a linear relationship between the two.

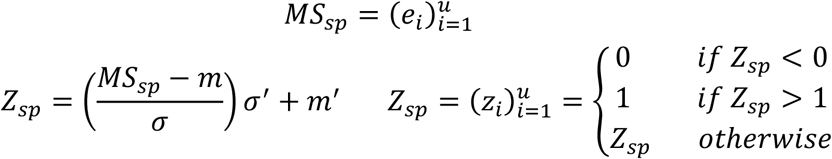

MS_sp_: module scores for the top 20 pseudotype markers calculated in spatial data, m, σ: mean and standard deviation of MS_sp_, m’, σ’: mean and standard deviation of the presumed pseudotype fraction, Z_sp_: presumed pseudotype fraction

To create a reference dataset, a pseudospot was aggregated to the pseudotype of a randomly chosen spot, and the combined gene expression for the composite marker panel was calculated (**Supplementary Fig. 12**). The resulting cell mixture was named the modified pseudospot. Since pseudotype markers were selected according to the log fold change in the pseudobulk approach, the expression of pseudotype markers in pseudotypes is expected to be significantly higher than that of single-cell markers. Additionally, for simplicity, the expression of all pseudotype markers in a spot was assumed to be directly proportional to that in the pseudotype of the spot. Thus, the single-cell marker expression in a modified pseudospot was set to 0, and the pseudotype marker expression was assigned by multiplying the normalized count of the selected spot by the presumed pseudotype fraction.

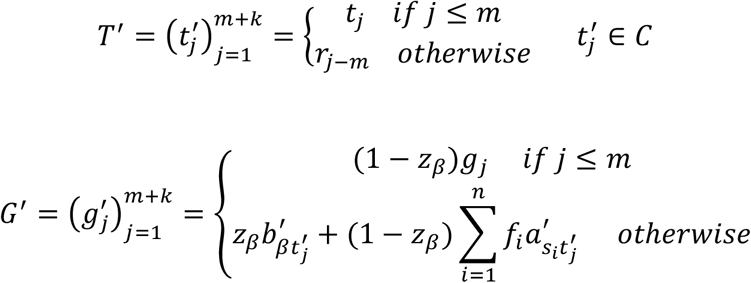

T’: index of the marker genes and pseudotype markers in the single-cell count matrix (RC_sc_), G’: composite gene expression profiles of the modified cell mixture (named modified pseudospot), β: randomly selected index of the spot

Finally, the gene expression profiles of the pseudotype and the pseudospot were summed with the pseudotype to nonpseudotype ratio as a weight, and the integrated expression of a modified pseudospot was obtained (**Supplementary Fig. 12**). Of note, the pseudotype fraction was extracted from the same spatial spot that the expression profile was referenced. The algorithm for generating a modified pseudospot was summarized with the above formulae. The modified pseudospots and spatial transcriptomic data were jointly provided as inputs for the domain adaptation in CellDART^9^.

#### Exploration of an optimal parameter range

The key parameters for spSeudoMap are the number of markers per single-cell cluster (*N_ markers*), the ratio of the total number of single-cell to pseudotype markers (*m/k* ratio), and the mean and standard deviation of the pseudotype fraction in spatial spots (*pseudof_m* and *pseudof_std*). The performance of the spSeudoMap was tested across the various parameters in human DLPFC datasets (slide number: 151676). Since the proportion of 10 layer-specific excitatory neurons was 0.53 among the single-cell data, *pseudof_m* was set to 0.47. The cortical layer annotation in spatial data was used as a reference, and a layer discriminative accuracy of the predicted neuronal fraction was assessed by an area under the receiver operating characteristic curve (AUROC). In general, spSeudoMap was capable of stably predicting the spatial distribution of neuron subpopulations with a median AUC over 0.5 with *N_markers* larger than 20, *m/k* larger than 1, and *pseudo_std* larger than 0.05 (**Supplementary Fig. 2**). The corresponding parameter ranges were selected for the downstream analyses. For the human brain (slide 151673) and mouse brain tissues, *N_markers* was set to 80, the *m/k* ratio to 4, and *pseudof_std* to 0.1. In human breast cancer tissue, *N_markers* was set to 40, the *m/k* ratio to 2, and *pseudof_std* to 0.1. Other parameters for the domain adaptation were given as the user guidelines of CellDART^9^.

#### Performance evaluation of spSeudoMap

The performance of spSeudoMap to predict the spatial cell composition of subpopulation single-cell data was evaluated in the three different datasets. For human DLPFC tissue, the accuracy of each model to localize 10 excitatory neuron types to corresponding layers was measured by AUROC. In the case of the mouse brain sample, the spot clusters were named after the anatomical region, and the spatial distribution of region-specific excitatory neuron types across the clusters was examined. Finally, in human breast tissue, the spatial localization patterns of the top immune cell subpopulation were mapped. The spatial correlation patterns were evaluated by Spearman’s correlation coefficient and visualized with a heatmap.

For the assessment of spSeudoMap, CellDART and Cell2location were selected as reference methods. The original single-cell dataset covering all cell types was jointly analyzed with a spatial transcriptome. CellDART deconvolutes spatial cell proportion based on domain adaptation. The numbers of sampled cells in the cell mixture were set to 8 and 10 in brain and breast cancer tissues, respectively. The number of markers per cell cluster was set to 20, and other parameters were given as the authors suggested. In addition, Cell2location^10^, which predicts spatial cell proportion based on the Bayesian statistical model, was selected for the comparison. In a previous study, it showed a similar performance to CellDART in spatially localizing layer-specific neuron types^9^. The expected cell abundance in each spatial spot was set to 8 in the brain tissue. The number of iterations was 30,000, and default values were given to the rest of the parameters in the model.

## Data availability

Publicly available transcriptomic datasets were utilized for the analyses. First, human DLPFC single-nucleus and spatial transcriptomic (slide no. 151673 and 151676) datasets were downloaded from http://research.libd.org/spatialLIBD/and_GSE144136, respectively. Second, mouse brain coronal section single-nucleus and spatial datasets were acquired from E-MTAB-11115 and 10X Genomics data repository, respectively. Lastly, matched single-cell and spatial data from the breast cancer patient were downloaded from GSE176078 and https://doi.org/10.5281/zenodo.3957257, while CD45+ sorted single-cell data from GSE114726.

## Code availability

Python source code and R wrap function for spSeudoMap is uploaded on https://github.com/bsungwoo/spS_eudoMap.

