## Supplementary Fig for "spSeudoMap: Cell type mapping of spatial transcriptomics using unmatched single-cell RNA-seq data"

**a**

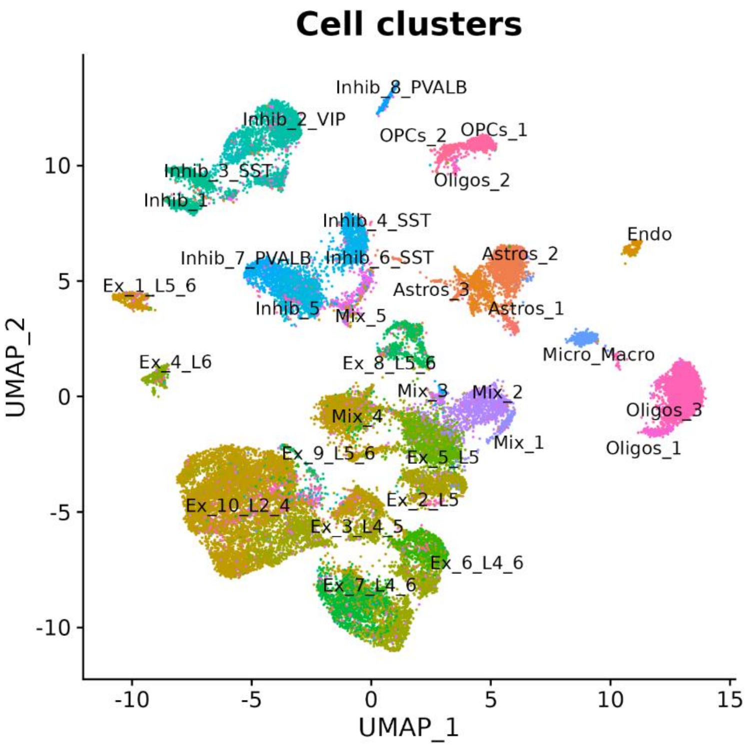
Cell clusters
Inhib_8_PVALB
Inhib_2_VIP
OPCs_2
OPCs_1
Inhib_3_SST
Oligos_2
Inhib_1
Inhib_4_SST
Endo
Inhib_7_PVALB
Inhib_6_SST
Astros_2
Ex_1_L5_6
Astros_3
Inhib_5
Mix_5
Astros_1
Micro_Macro
Ex_4_L6
Ex_8_L5_6
Mix_3
Mix_2
Mix_4
Oligos_3
Mix_1
Oligos_1
Ex_9_L5_6
Ex_5_L5
Ex_10_L2_4
Ex_2_L5
Ex_3_L4_5
Ex_6_L4_6
Ex_7_L4_6
UMAP_2
UMAP_1

**b**

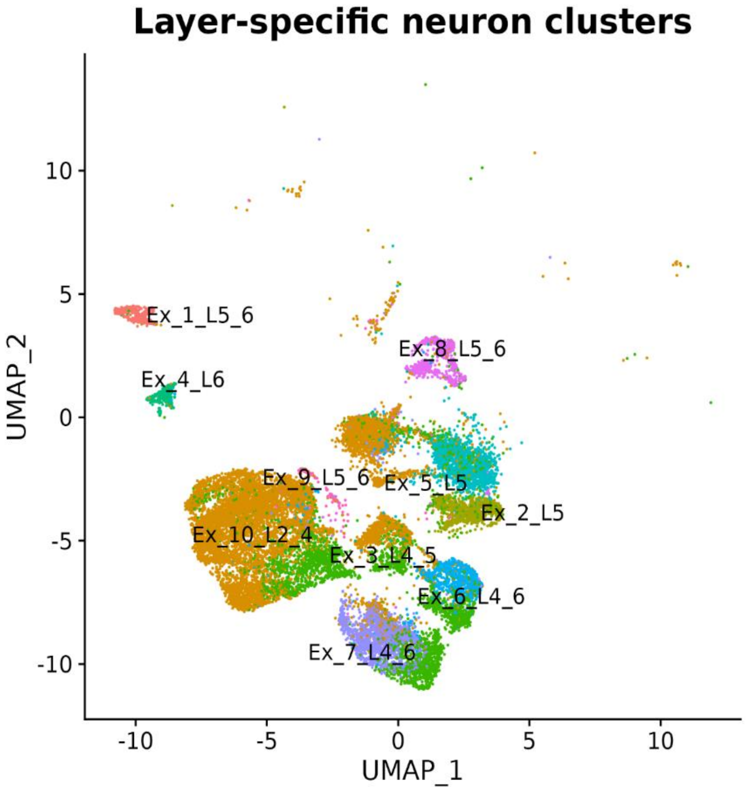
Layer-specific neuron clusters
Ex_1_L5_6
Ex_8_L5_6
Ex_4_L6
Ex_9_L5_6
Ex_5_L5
Ex_2_L5
Ex_10_L2_4
Ex_3_L4_5
Ex_6_L4_6
Ex_7_L4_6
UMAP_2
UMAP_1

**Supplementary Fig. 1. Single-nucleus datasets for the human DLPFC tissue**

**a,** Dimensionality reduction was performed for the single-nucleus data and the cells were visualized on a uniform manifold approximation and projection (UMAP) plot based on their expression profiles. The cell type annotation was color-coded on the plot.

**b,** Among the 33 cell types composing the brain, 10 layer-specific excitatory neurons were selected and utilized as the simulation dataset for the sorted single-cell data. The selected neuron types were visualized on UMAP.

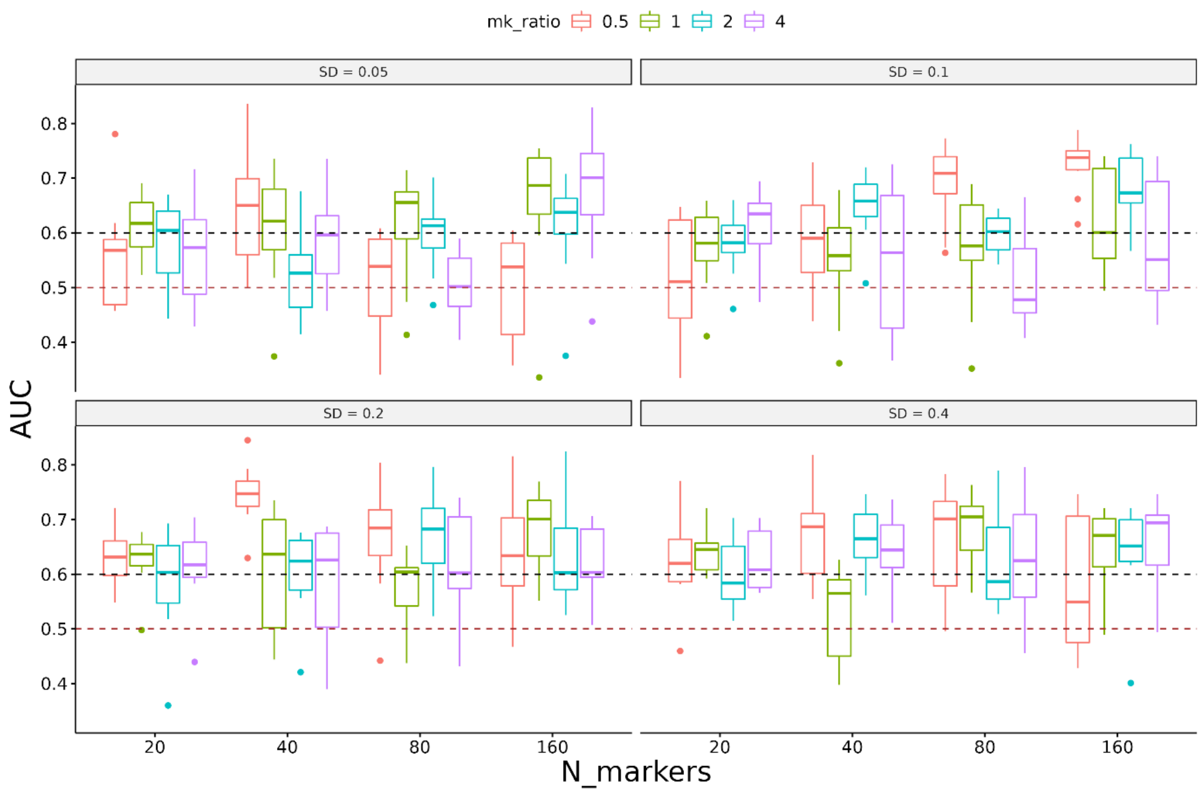


**Supplementary Fig. 2. Exploration of optimal parameters for spSeudoMap**

The optimal parameter ranges for spSeudoMap were searched in human brain tissue (slide number: 151676). The main parameters composing the model, the number of markers (*N_markers*), ratio of the number of single-cell markers to pseudotype markers (*m/k* ratio), and standard deviation of pseudotype fraction (SD) were changed and the performance was evaluated. The 10 layer-specific excitatory neuron fraction predicted by spSeudoMap was assessed whether it precisely localizes the cell type to the corresponding cortical layer. Receiver operating characteristic analysis was implemented and the area under the curve (AUC) was calculated for each cell type. The AUCs across 10 neuron types were pooled and visualized with boxplots. The plot revealed that the performance of spSeudoMap was overall stable when *N_markers*>20, *m*/*k* ratio>1, and *pseudo_std*> 0.05.

**a** **CellDART with subpopulation**

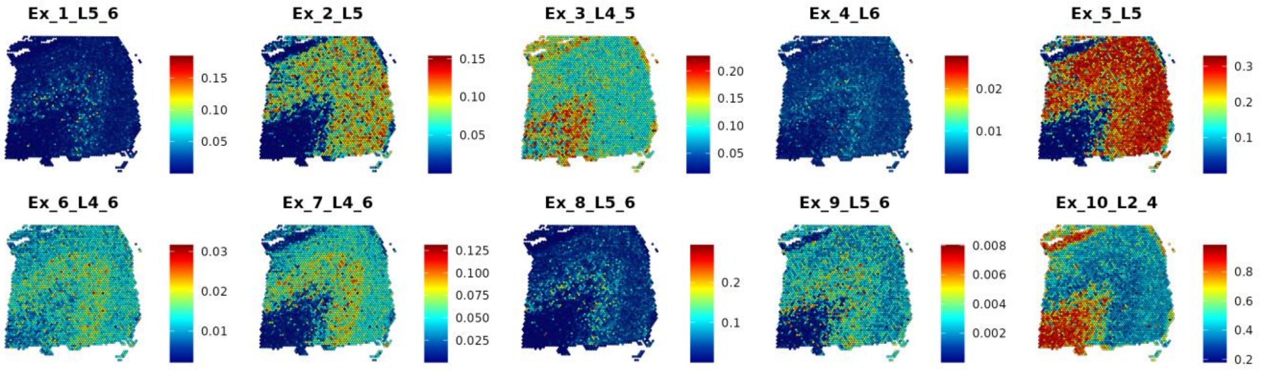


**b** **Cell2location with subpopulation**

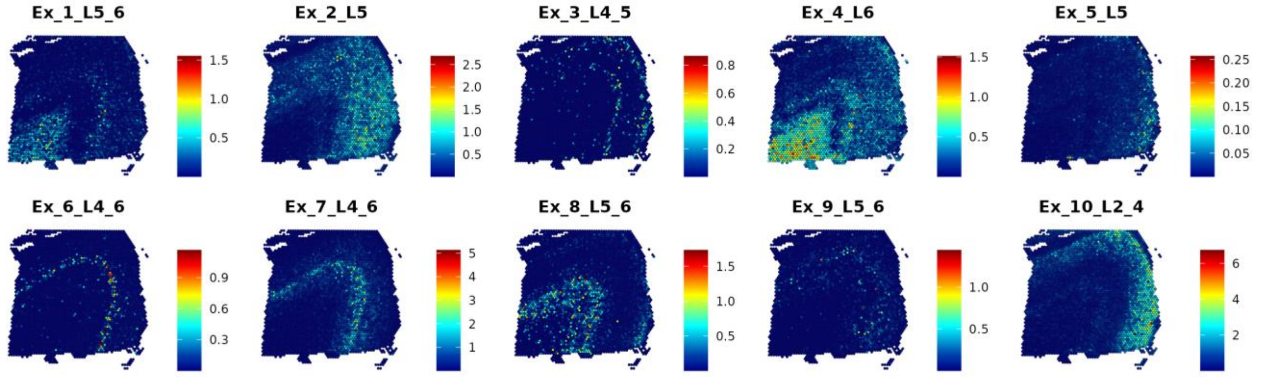


**Supplementary Fig. 3. Spatial distribution patterns of layer-specific neurons estimated by CellDART and Cell2location**

The spatial composition of 10 layer-specific neuron types in human DLPFC tissue (slide number: 151673) was predicted by integrating spatial data with simulated subpopulation single-cell data. It was to assess the performance of the existing models, CellDART and Cell2location, in spatially mapping cell subpopulations. **(a)** In CellDART, the neuron types were localized to a corresponding cortical layer in the majority of the cases; however, it failed to reflect the layer-specificity in Ex_3_L4_5 and Ex_10_L2_4. **(b)** In Cell2location, the cell types were highly restricted to the expected cortical layers except for Ex_1_L5_6 and Ex_9_L5_6.

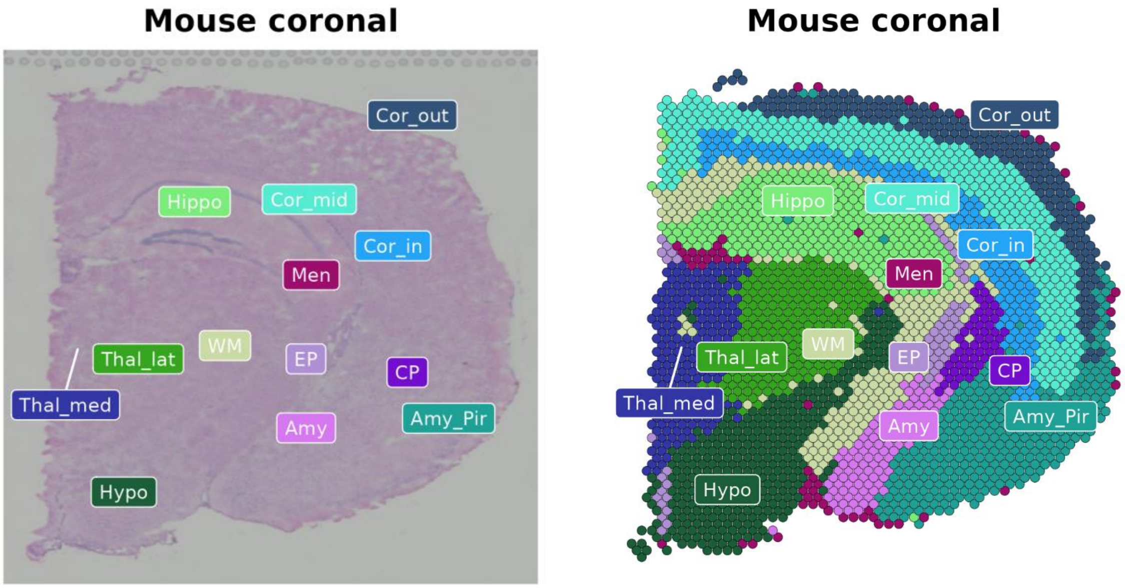


**Supplementary Fig. 4. Spot clustering of mouse brain spatial transcriptome**

The spots comprising the spatial data were clustered based on the expression of highly variable genes (HVGs). Then, the resulting clusters were renamed according to the anatomical locations and visualized on top of the tissue. Amy: amygdala, Amy_Pir: amygdala or piriform cortex, Cor_out: outer cortex, Cor_mid: mid cortex, Cor_in: inner cortex, CP: caudoputamen, EP: ependyma, Hippo: hippocampus, Hypo: hypothalamus, Men: meninges, Thal_lat: lateral thalamus, Thal_med: medial thalamus, and WM: white matter.

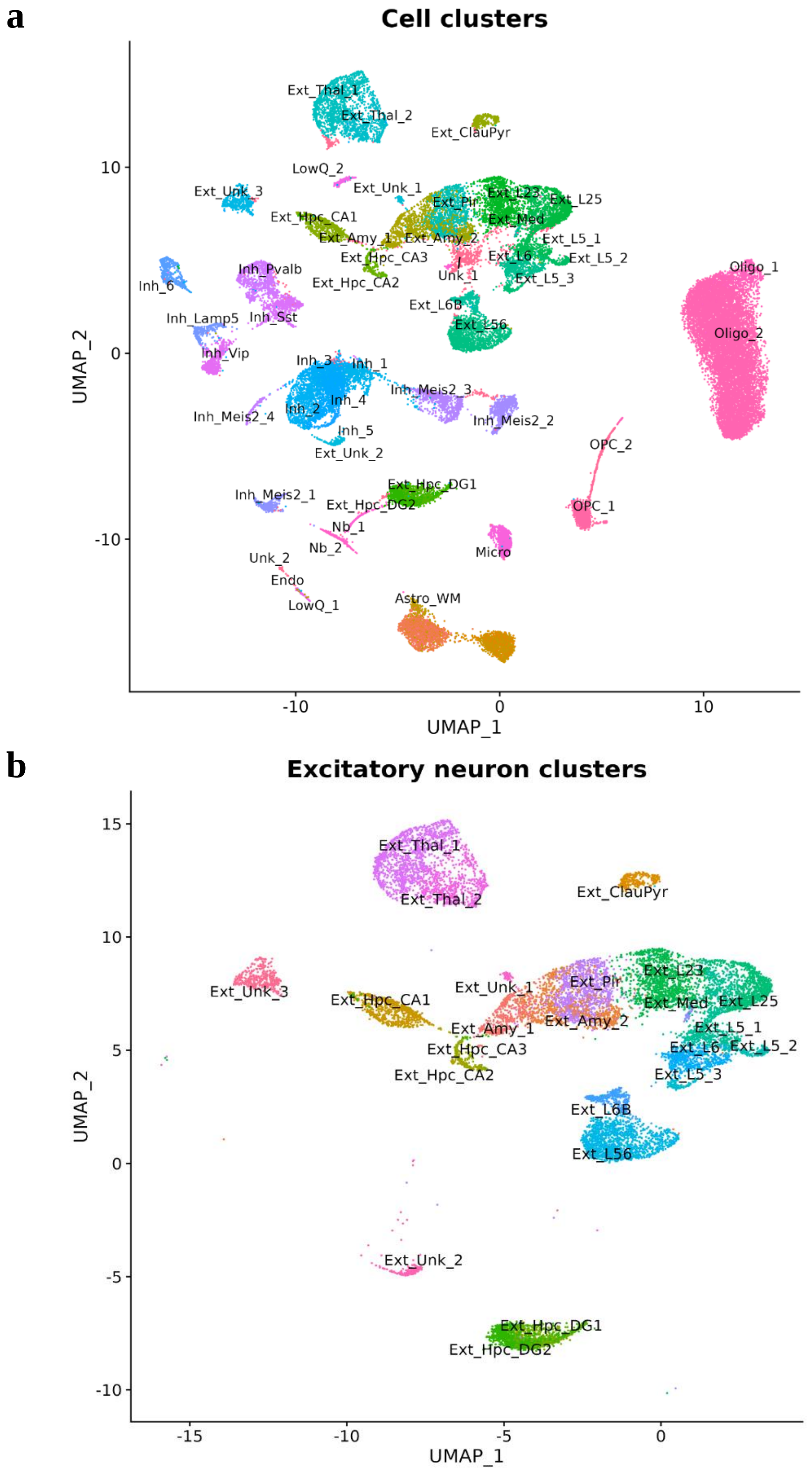
a
Cell clusters
UMAP_1
UMAP_2
Ext_Thal_1
Ext_Thal_2
Ext_ClauPyr
LowQ_2
Ext_Unk_3
Ext_Unk_1
Ext_Hpc_CA1
Ext_Pir
Ext_L23
Ext_L25
Ext_Med
Ext_Amy_1
Ext_Amy_2
Ext_L5_1
Ext_Hpc_CA3
Ext_L6
Ext_L5_2
Ext_Hpc_CA2
Unk_1
Ext_L5_3
Inh_6
Inh_Pvalb
Oligo_1
Ext_L6B
Inh_Lamp5
Inh_Sst
Ext_L56
Oligo_2
Inh_Vip
Inh_3
Inh_1
Inh_Meis2_3
Inh_4
Inh_Meis2_4
Inh_2
Inh_Meis2_2
Inh_5
Ext_Unk_2
OPC_2
Inh_Meis2_1
Ext_Hpc_DG1
Ext_Hpc_DG2
OPC_1
Nb_1
Nb_2
Micro
Unk_2
Endo
LowQ_1
Astro_WM
b
Excitatory neuron clusters
UMAP_1
UMAP_2
Ext_Thal_1
Ext_Thal_2
Ext_ClauPyr
Ext_Unk_3
Ext_Hpc_CA1
Ext_Unk_1
Ext_Pir
Ext_L23
Ext_Med
Ext_L25
Ext_Amy_1
Ext_Amy_2
Ext_L5_1
Ext_Hpc_CA3
Ext_L6
Ext_L5_2
Ext_Hpc_CA2
Ext_L5_3
Ext_L6B
Ext_L56
Ext_Unk_2
Ext_Hpc_DG1
Ext_Hpc_DG2

**Supplementary Fig. 5. Single-nucleus data for the mouse brain coronal section**

**a,** Dimensionality reduction was performed and the cells composing single-nucleus data were visualized on a UMAP plot. The cell type annotation was color-coded on the plot.

**b,** The 23 region-specific excitatory neurons were selected from a total of 59 cell types in the brain. The opted neuron types were visualized on UMAP and were utilized as a simulation dataset for the sorted single-cell data.

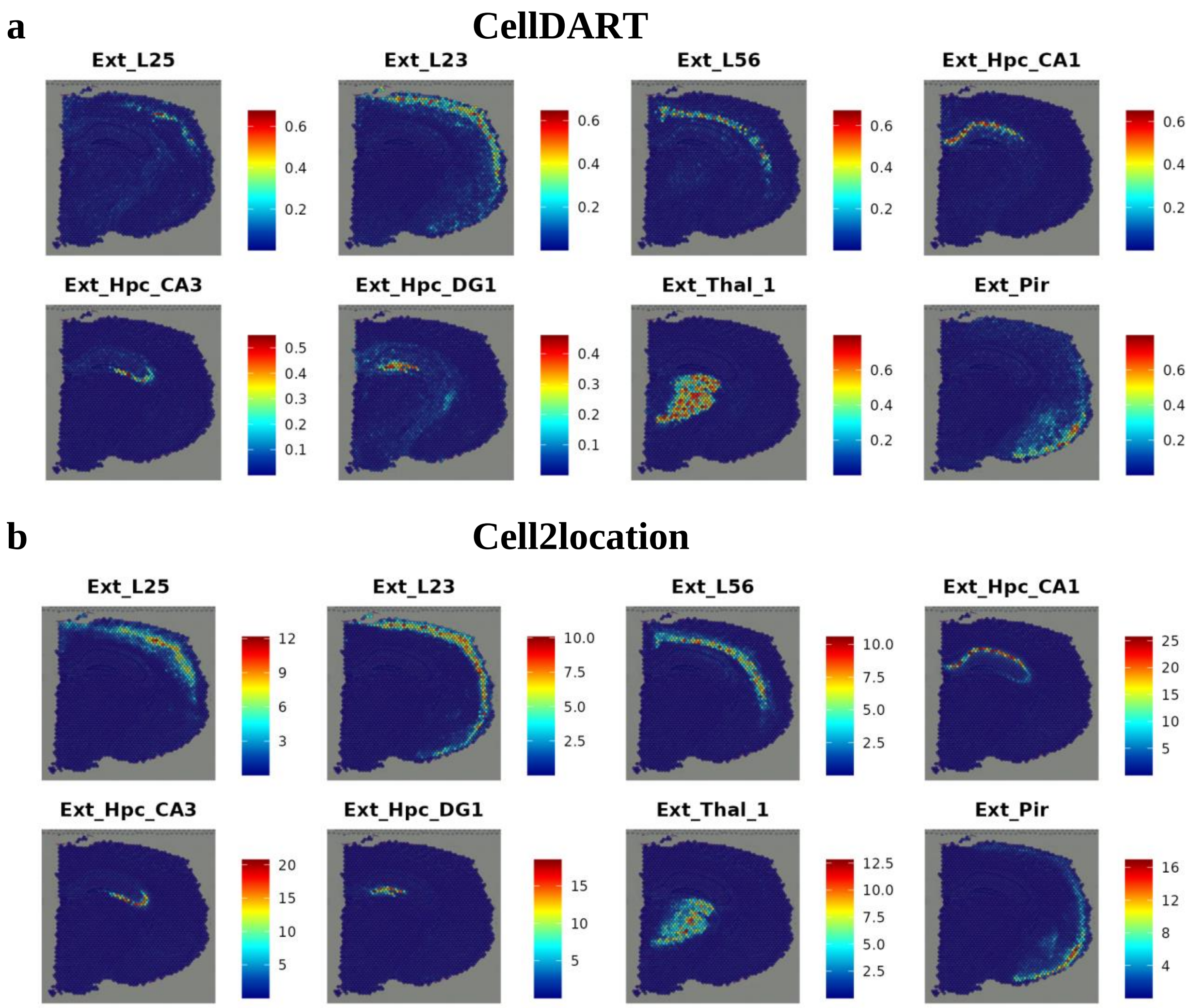


**Supplementary Fig. 6. Spatial maps of region-specific neurons in mouse brain using original single-cell data containing all cell types: the representative neuron types**

The representative excitatory neuron types were spatially mapped to the tissue using original single-cell data containing all cell types. The spatial maps from the two methods, CellDART and Cell2location, were considered references to evaluate the prediction results of spSeudoMap. For both **(a)** CellDART and **(b)** Cell2location, the neuron types were spatially localized to the expected anatomical regions: Ext_L25, Ext_L23, and Ext_L56 to corresponding cortical layers, Ext_Hpc_CA1, Ext_Hpc_CA3, and Ext_Hpc_DG1 to the hippocampus, Ext_Thal_1 to the thalamus, and Ext_Pir to piriform cortex.

**a** **CellDART with all cell types**

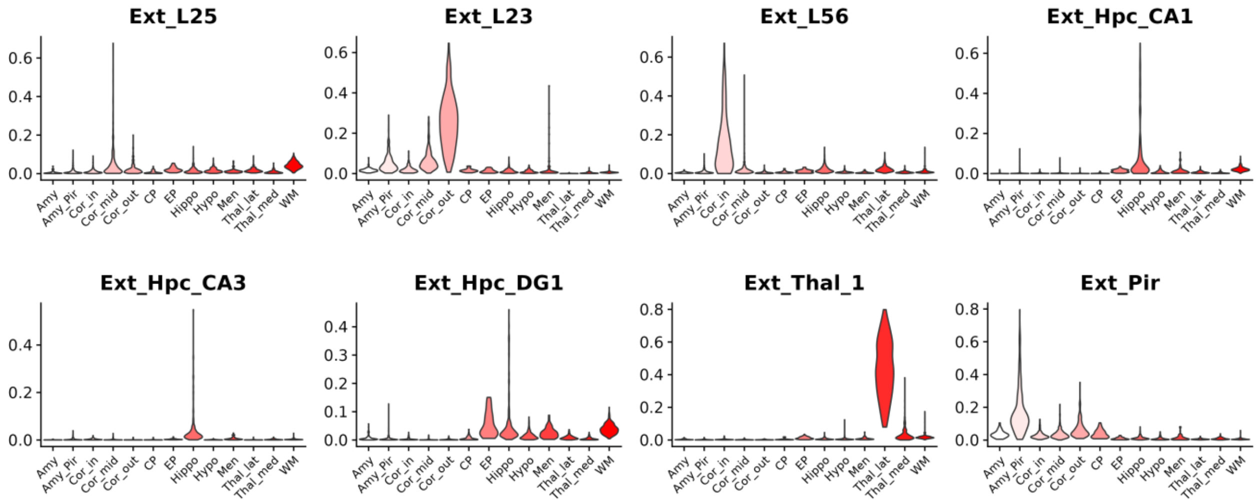


**b** **Cell2location with all cell types**

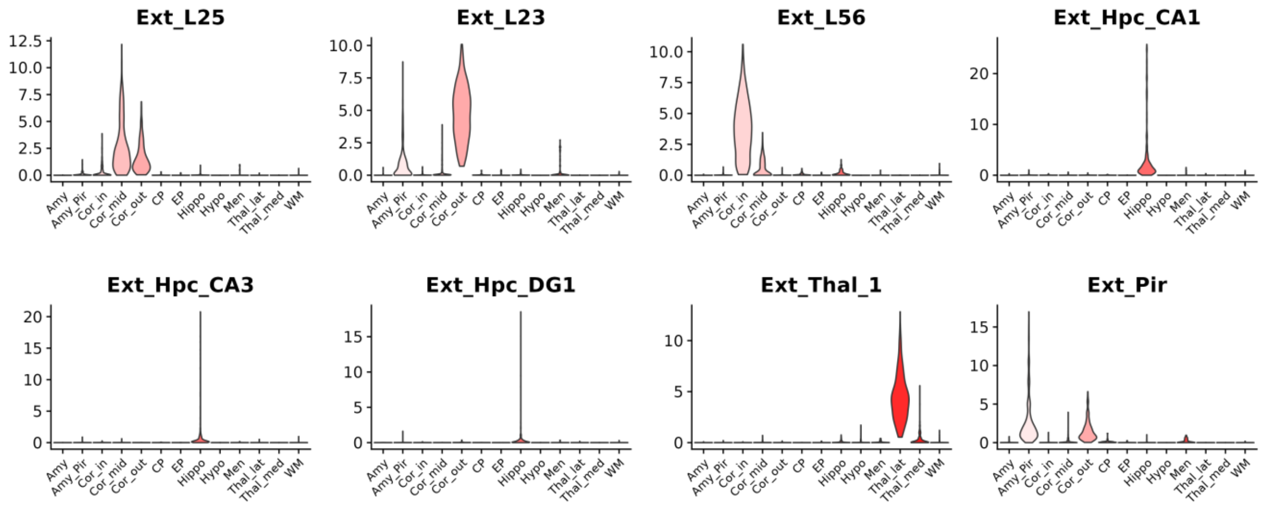

**Supplementary Fig. 7. Distribution of the neuron subtypes in mouse brain across the locations predicted from single-cell data covering all cell types**

The region-specific excitatory neuron types were mapped to the tissue by CellDART and Cell2location. Violin plots represent the predicted excitatory neuron fraction in the corresponding spot clusters. In both **(a)** CellDART and **(b)** Cell2location, the cell subtypes were accurately localized to the expected anatomical locations. Amy: amygdala, Amy_Pir: amygdala or piriform cortex, Cor_out: outer cortex, Cor_mid: mid cortex, Cor_in: inner cortex, CP: caudoputamen, EP: ependyma, Hippo: hippocampus, Hypo: hypothalamus, Men: meninges, Thal_lat: lateral thalamus, Thal_med: medial thalamus, and WM: white matter.

**a** **CellDART with all cell types**

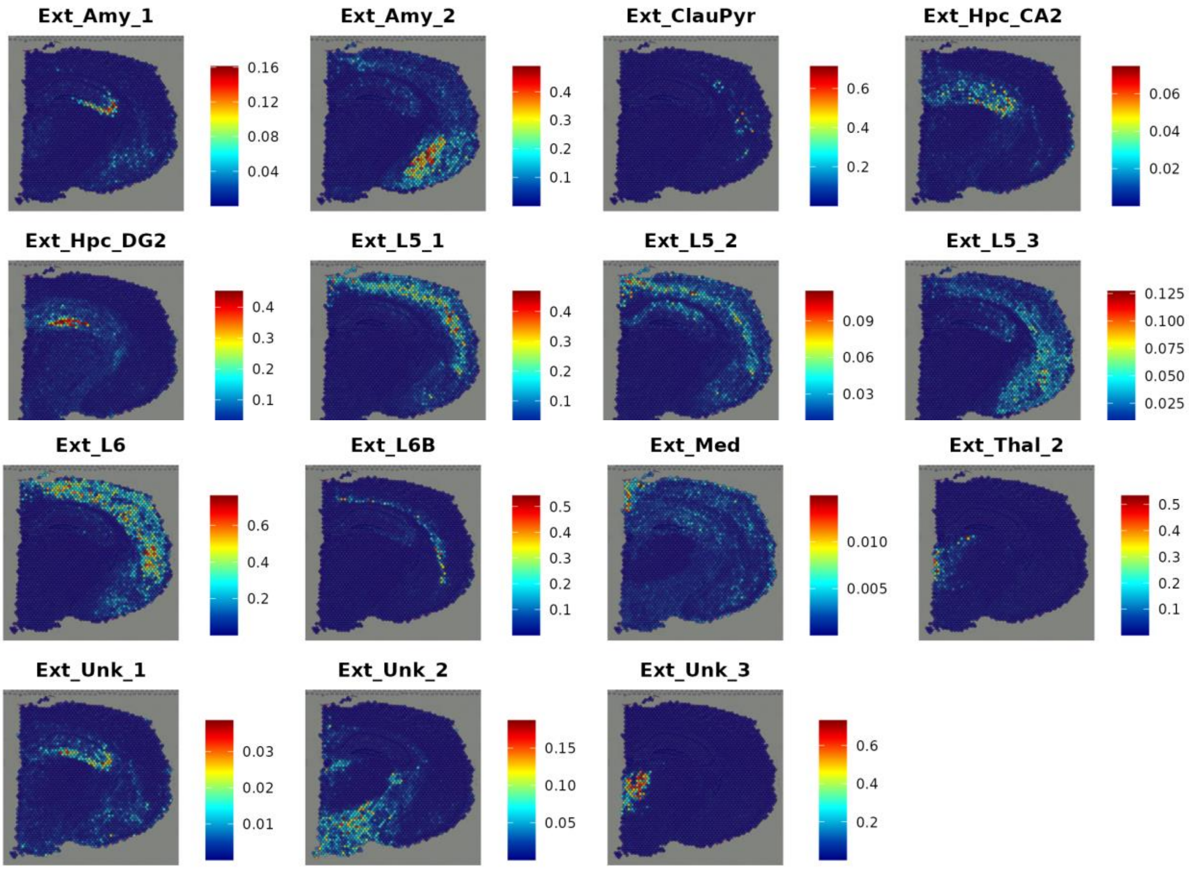

**b** **Cell2location with all cell types**

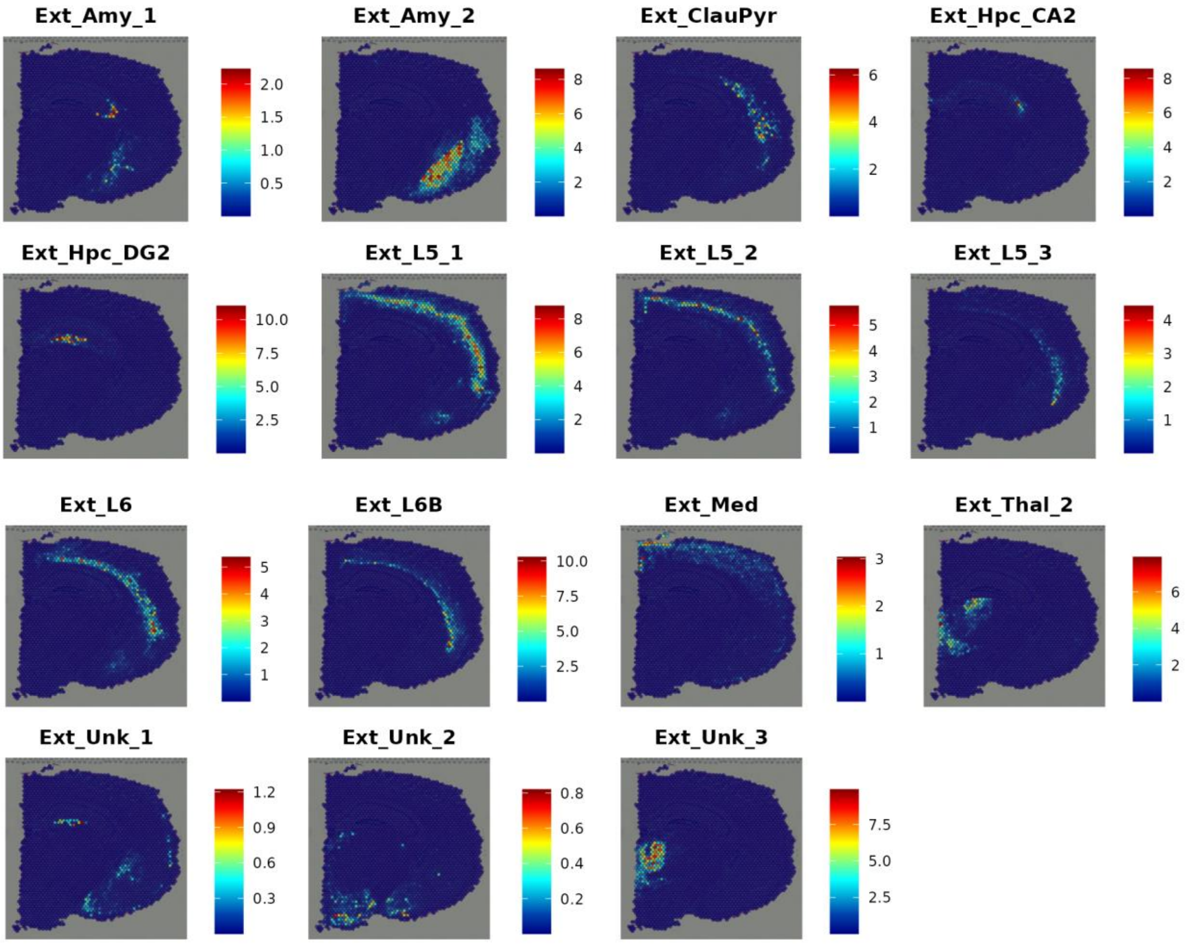

c

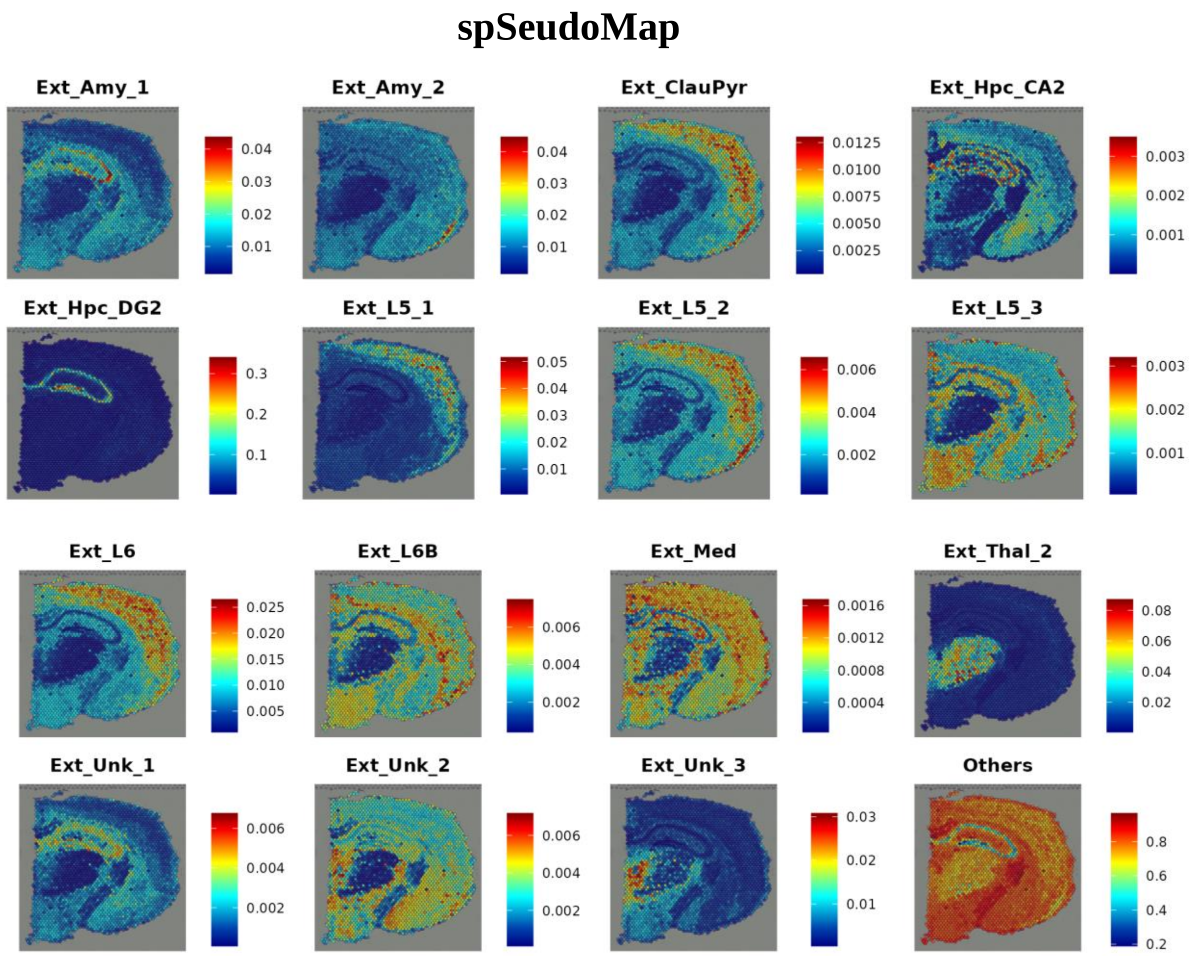
spSeudoMap
Ext_Amy_1
Ext_Amy_2
Ext_ClauPyr
Ext_Hpc_CA2
Ext_Hpc_DG2
Ext_L5_1
Ext_L5_2
Ext_L5_3
Ext_L6
Ext_L6B
Ext_Med
Ext_Thal_2
Ext_Unk_1
Ext_Unk_2
Ext_Unk_3
Others

**Supplementary Fig. 8. The spatial landscape of region-specific neurons in mouse brain: rest of the neuron types**

The spatial distribution of the rest of the region-specific neuron types in the mouse brain was predicted by integrating spatial data with single-cell data covering all cell types using **(a)** CellDART and **(b)** Cell2location. The spatial maps created by both tools were considered references and were compared with the cell subpopulation mapping results from **(c)** spSeudoMap. In both **(a)** CellDART and **(b)** Cell2location, the layer-specific neurons were localized to the corresponding cortical layers, hippocampal neurons to the hippocampus, and thalamic neurons to the thalamus (except for Ext_Unk neuron types whose regional specificity is not known). Overall the two reference methods showed similar patterns of spatial distribution. **(c)** In the case of spSeudoMap, the rest of the neuron types showed high cellular fractions in the corresponding anatomical locations (except for Ext_Unk neuron types whose regional specificity is not known). However, the proportion was also high in non-specific regions, particularly in the cell types with low cell fractions.

a

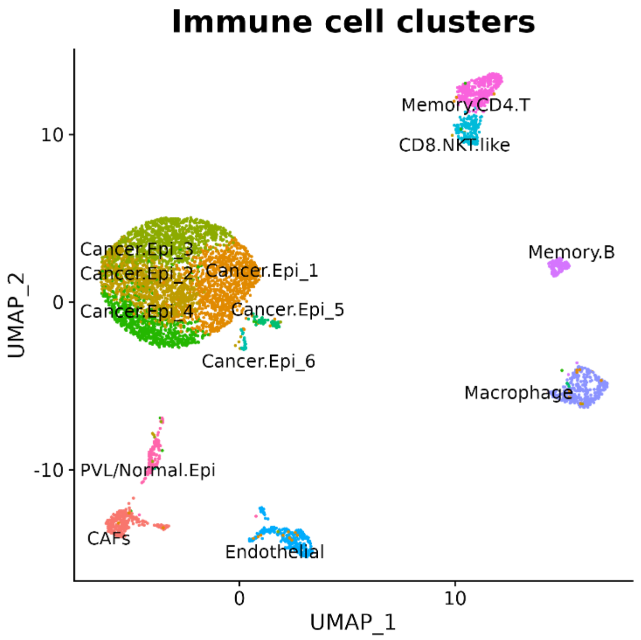
Immune cell clusters
Memory.CD4.T
CD8.NKT.like
Memory.B
Cancer.Epi_3
Cancer.Epi_2
Cancer.Epi_1
Cancer.Epi_4
Cancer.Epi_5
Cancer.Epi_6
Macrophage
PVL/Normal.Epi
CAFs
Endothelial
UMAP_1
UMAP_2

b

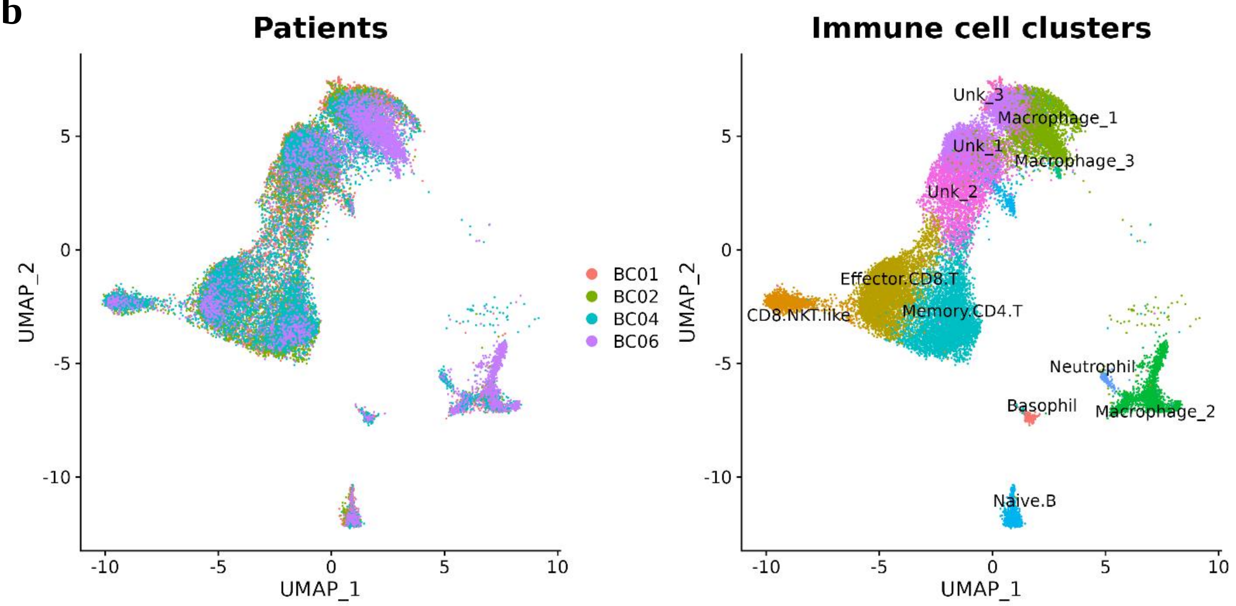
Patients
BC01
BC02
BC04
BC06
Immune cell clusters
Unk_3
Macrophage_1
Unk_1
Macrophage_3
Unk_2
Effector.CD8.T
CD8.NKT.like
Memory.CD4.T
Neutrophil
Basophil
Macrophage_2
Naive.B
UMAP_1
UMAP_2

**Supplementary Fig. 9. Single-cell data for the human breast cancer**

**a,** Dimensionality reduction was performed for the matched single-cell data of the same patient. The gene expression profile of cells was visualized with a UMAP plot and the cell types were color-coded. The dataset was considered as a reference for the evaluation of spatial cell composition.

**b,** Dimensionality reduction was done and the single-cell data obtained by CD45+ sorting was exhibited using a UMAP plot. Only the 12 immune cell types were selected and visualized. Then the immune cell types were spatially mapped to the tissue using spSeudoMap.

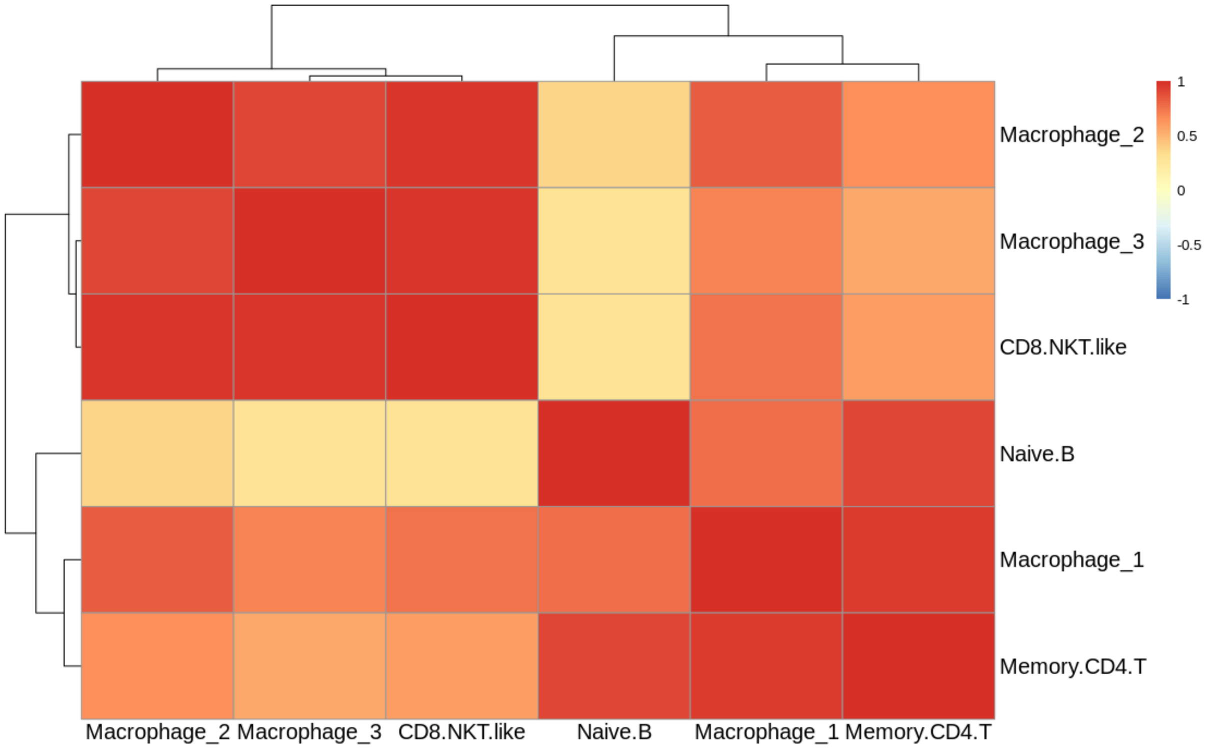


**Supplementary Fig. 10. Spatial correlation patterns between immune cells in the human breast cancer tissue**

Spearman's correlation coefficients were calculated between the predicted cell fraction of the top 8 cell types. The proximity of the spatial correlation patterns was examined and hierarchical clustering was performed. Memory CD4 T cells and macrophage_1, and CD8+ NKT-like cells and macrophage_3 showed high similarities among the cell type pairs.

**a**

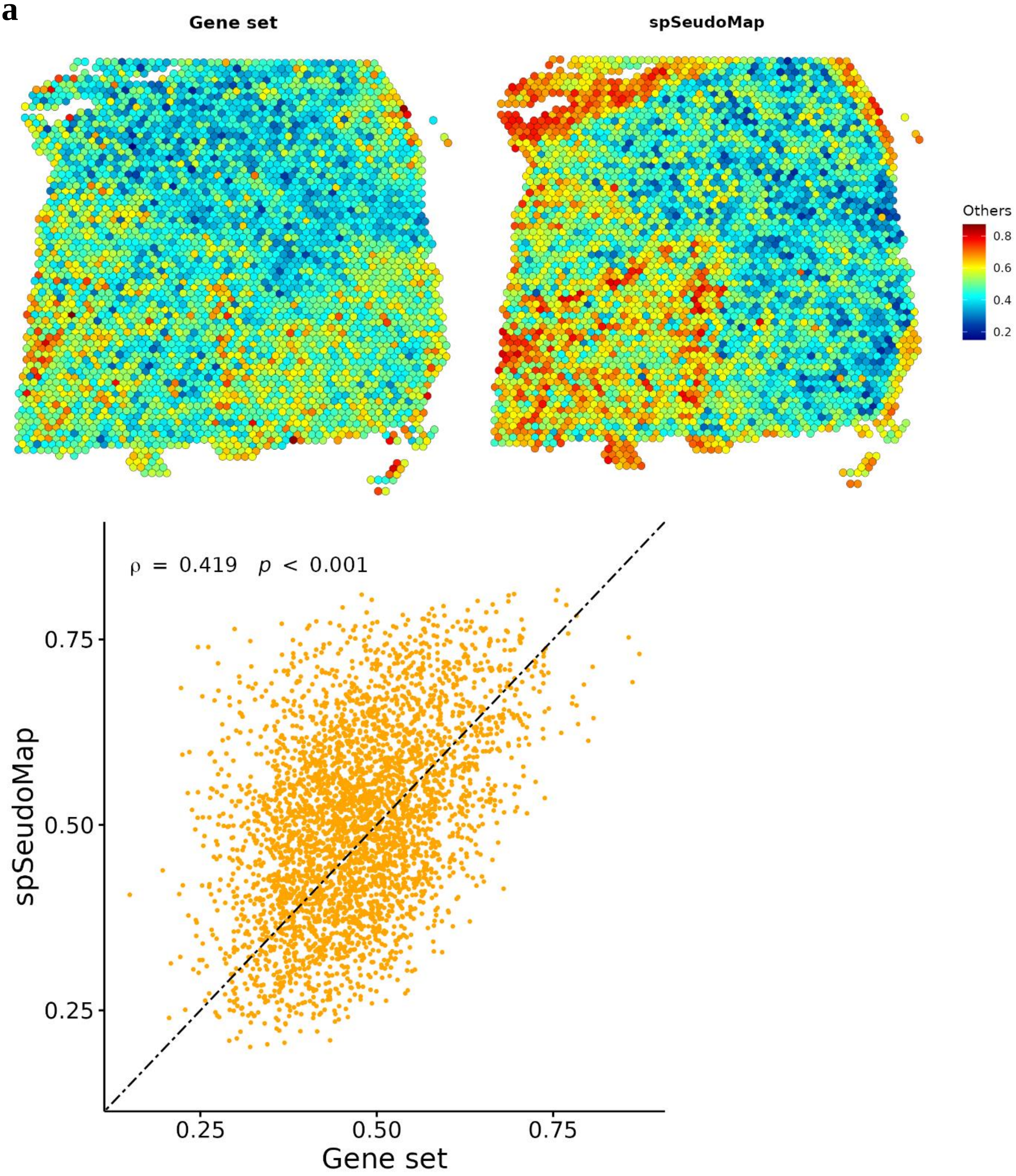

Gene set
spSeudoMap
Others
0.8
0.6
0.4
0.2
ρ = 0.419 p < 0.001
0.75
0.50
0.25
spSeudoMap
0.25
0.50
0.75
Gene set

**b**

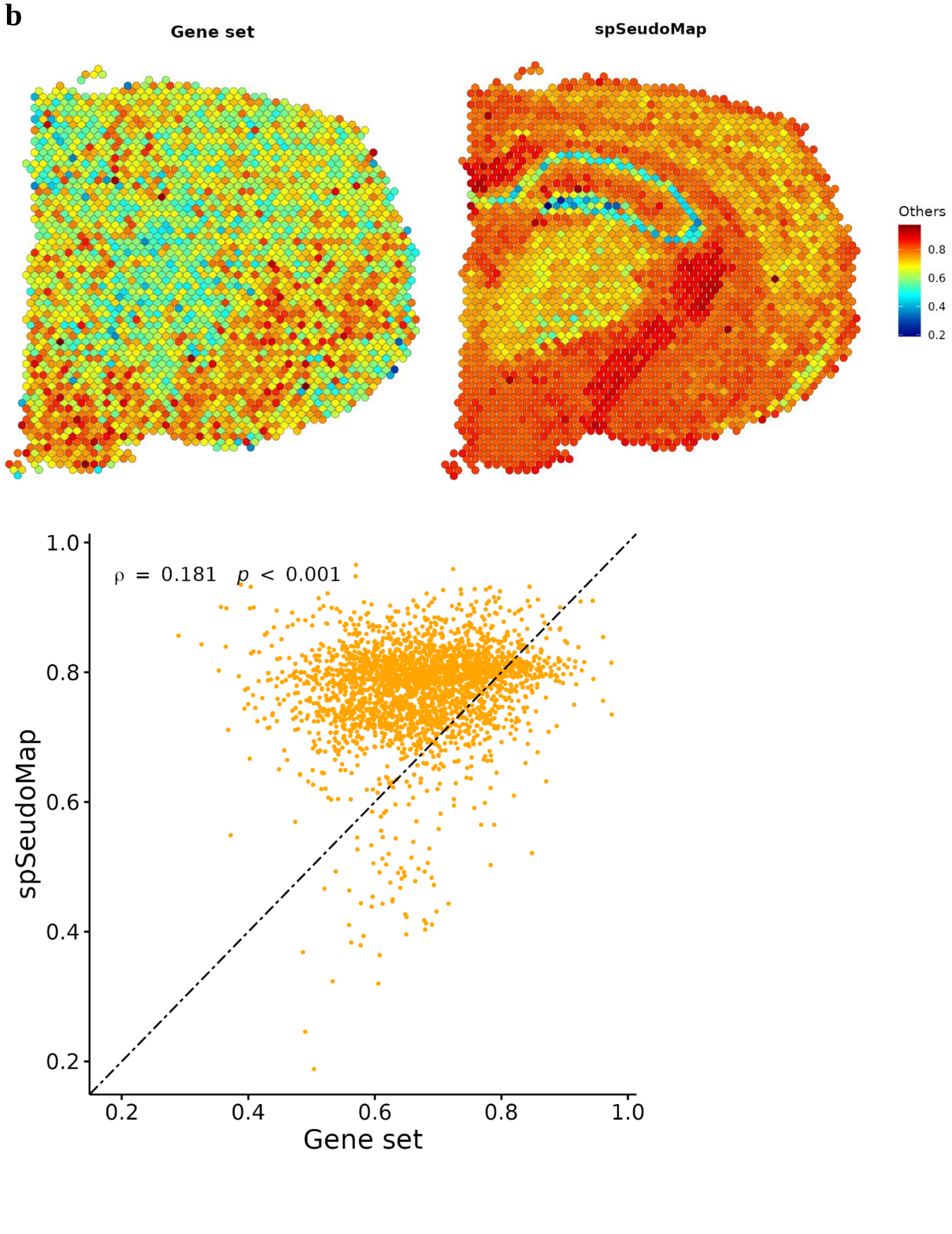

Gene set
spSeudoMap
Others
0.8
0.6
0.4
0.2
ρ = 0.181 p < 0.001
1.0
0.8
0.6
0.4
0.2
spSeudoMap
0.2
0.4
0.6
0.8
1.0
Gene set

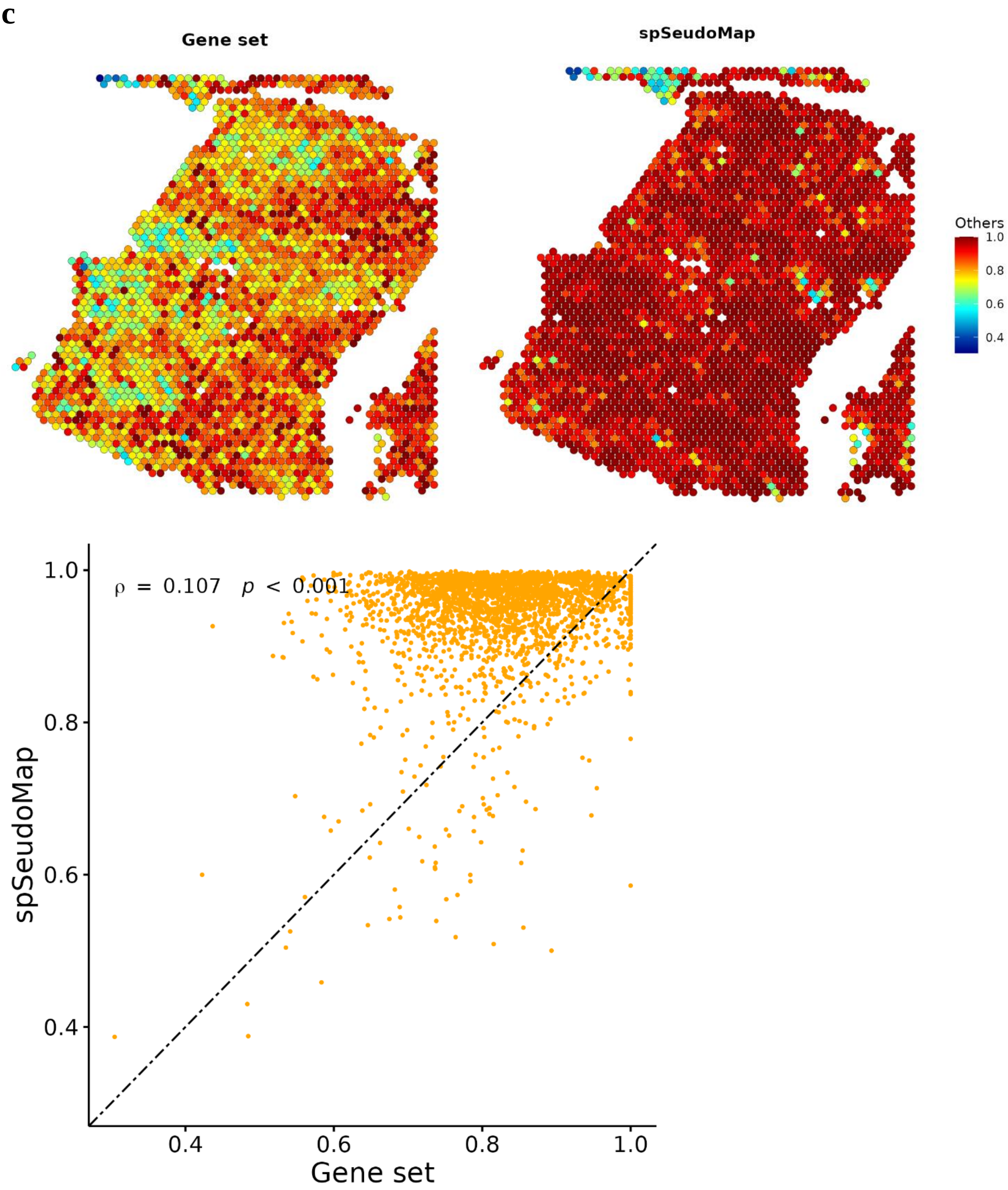
c
Gene set
spSeudoMap
Others
1.0
0.8
0.6
0.4
ρ = 0.107 p < 0.001
spSeudoMap
Gene set
0.4
0.6
0.8
1.0

**Supplementary Fig. 11. Comparison between pseudotype fraction used for modified pseudospot generation and that calculated from spSeudoMap.**

The two different estimation results for pseudotype fraction in each spatial spot were mapped to the tissue and scatter plots were shown. In the top-left panel and the x-axis of the scatter plots, the pseudotype fraction calculated from the gene set scores of the top 20 pseudotype markers was presented. In the top-right panel and the y-axis of the scatter plots, pseudotype fraction predicted as the output of spSeudoMap was exhibited. Spearman's correlation coefficients and statistical significance were calculated and presented in the top-left corner of each plot. The two fraction values showed positive correlations in **(a)** human brain (slide number: 151673), **(b)** mouse brain, and **(c)** human breast cancer tissues.

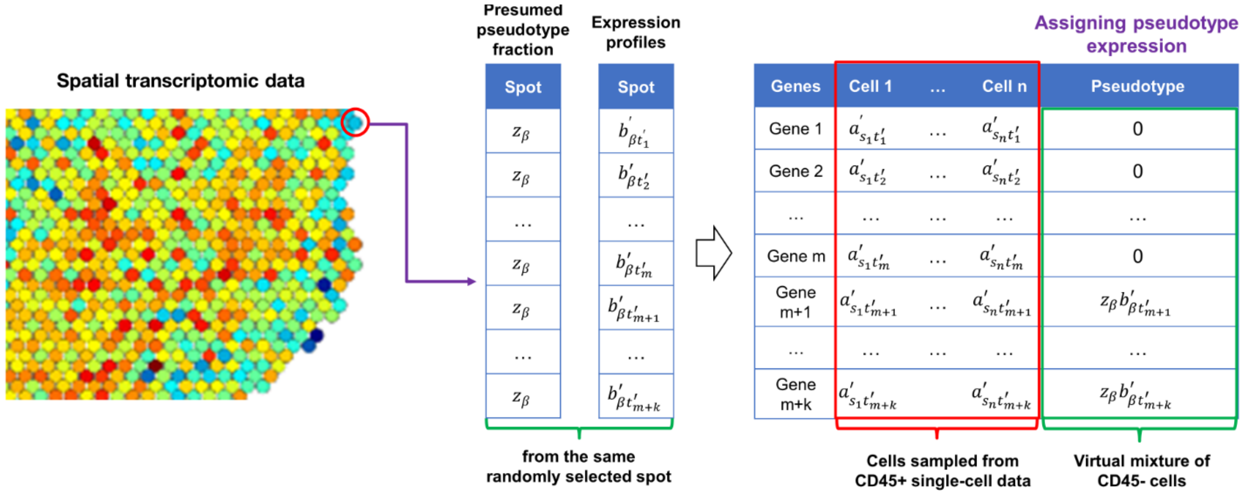


**Supplementary Fig. 12. The schematic diagram for creating modified pseudospots, the reference dataset for the domain adaptation**

A spatial spot was randomly sampled from the spatial transcriptomic data. The presumed pseudotype fraction ($z_\beta$) and expression profiles of the spot ($b'_{\beta t'_1}$, $b'_{\beta t'_2}$, $\cdots$ $b'_{\beta t'_{m+k}}$) were referenced. Then, the expression profiles of virtual pseudotype markers (Gene m+1, ⋯ Gene m+k) in pseudotype were assigned by multiplying the expression profiles of the selected spot with the pseudotype fraction ($z_\beta b'_{\beta t'_j}$). The expression of the rest of the maker genes (Gene 1, ⋯ Gene m) in pseudotype was set to 0. Finally, to create a modified pseudospot, a cell mixture that closely resembles the spatial spots, pseudotype expression was aggregated with composite gene expression profiles of a cell mixture generated from a random sampling of cells from the single-cell data (e.g. CD45+ sorted single-cell RNA sequencing data).
